# Tunable bet hedging in yeast responses to osmotic stress

**DOI:** 10.1101/039982

**Authors:** Yoshikazu Hirate, Samuel Bottani, Wyming Lee Pang, Suzannah Rutherford

## Abstract

Microbes limit risk by stochastic bet hedging – low frequency expression of less fit, slow growing cells constitutively preadapted against many stresses including antibiotics. By contrast, here we report continuous variation in the *induced* frequency of cells with slow osmotic stress signaling, survival and proliferation among 50 ecologically-distinct strains of budding yeast challenged by sudden hyperosmotic stress. Despite extensive variation in early mortality, strains displayed robust perfect adaptation and recovery of steady-state viability in moderate stress. In severe stress survival depended on strain-specific proportions of cells with divergent strategies. ‘Cautious’ cells survived without dividing; ‘reckless’ cells attempted to divide too soon and failed, killing both mother and daughter. We show that heritable frequencies of cautious and reckless cells produce a rapidly diversifying template for microbial bet hedging that mimics natural variation in stress responses whose timing, amplitude and frequency could evolve – be ‘tuned’ by – different patterns of environmental stress.

---

An ability to sense and respond appropriately to environmental challenge can mean the difference between life and death. Organisms adapt physiologically to short-term stresses and evolve to track longer-term environmental change. Environmental stress responses present since life’s common ancestor are exquisitely adapted by natural selection in a diversity of habitats (Tomanek, 2010). For example heat shock proteins that allow survival of what would otherwise be killing temperatures are induced by slight elevations above optimal temperatures in thermal environments ranging from Antarctic to superheated hydrothermal vents, and have even evolved to anticipate and track predictable temperature fluctuations for organisms living in tide-pools (Feder and Hoffmann, 1999; Richter et al., 2010). By contrast, unpredictable and severe environmental stresses impose a higher-level selection over populations spanning multiple generations. Particularly for genetically related lineages or clones of microorganisms, genotypes that consistently produce a small fraction of cells with unsolicited stress responses (bet hedgers) are favored over genotypes that produce only optimally-fit individuals (Arnoldini et al., 2012; Donaldson-Matasci et al., 2013; King and Masel, 2007; Meyers and Bull, 2002). A trade-off between the growth arrest and cost of mounting unsolicited stress responses in preadapted individuals versus the potential escape from extinction for a diversified population or clone in unexpected and sudden stress favors the short term sacrifice of arithmetic mean fitness for longer-term multiplicative fitness (Arnoldini et al., 2012; De Jong et al., 2011; King and Masel, 2007; Simons, 2011). While pre-adaptive, stochastic stress resistance is a well-studied form of bet hedging in microorganisms, less is known about evolutionary forces governing physiological adaptation and the dangerous resumption of cell divisions in on-going episodes of environmental stress whose severity and duration are also often unpredictable.

The yeast high osmolarity glycerol (HOG) signaling pathway is central to an elaborate stress response that reduces cellular damage and death in unpredictable osmotic environments where the balance between external solutes and free water pressure in the cell can change suddenly (Hohmann, 2002). The HOG pathway consists of at least two highly-conserved, multi-component osmotic stress sensors linked to a parallel series of at least 15 kinases and accessory proteins that ultimately alter the activity of nearly 10% of the yeast genome (Hohmann, 2002; Saito and Posas, 2012). The sheer numbers of genes involved in HOG signaling, their conservation, and their elaborate circuitry suggest that a nuanced response to osmotic stress has been crucial and strongly selected throughout evolutionary history. However a main function of the HOG pathway is the production and accumulation of intracellular glycerol, which restores water balance and, as demonstrated by a large body of work from many labs, is essential for survival, physiological adaptation and proliferation in on-going hyperosmotic stress (Babazadeh et al., 2014; Clotet and Posas, 2007; Hohmann, 2002; Hohmann et al., 2007; Nadal et al., 2002; Saito and Posas, 2012). In the wild, yeast must balance immediate, individual survival against population-level evolutionary fitness. Individual fitness requires that cells carefully sense the amplitude and direction of environmental change and safely reenter the cell cycle after stress (Clotet and Posas, 2007). On the other hand, multiplicative fitness favors clonal populations that respond as rapidly as possible to improved conditions with on average earlier cell cycle reentry and faster proliferation – even if some individuals that reenter the cell cycle too quickly are lost (Ratcliff et al., 2014).

The hyperosmotic stress response of budding yeast is almost certainly under strong selection in nature and has well-characterized and accessible signaling and phenotypic traits that can be measured in the lab, making this an ideal system is ideal for characterizing the mapping between signaling behavior and fitness (Clotet and Posas, 2007; Hohmann, 2002; Saito and Posas, 2012). For example, glycerol-3-phosphate dehydrogenase (*GPD1*) is rate-limiting for glycerol production (Remize et al., 2001). We use the synthesis and accumulation of green fluorescent protein (GFP) integrated into the gene for *GPD1* as a proxy for HOG pathway activity (*GPD1*::GFP). To our knowledge bet hedging and developmental noise have been exclusively studied among cells or micro-colonies of a single or few strain backgrounds. Here we characterize natural variation in osmotic stress signaling, survival and adaptation in both exponentially growing and nearly quiescent cultures of diploid yeast. To that end we used a synthetic population of diverse yeast genotypes made by crossing *GPD1*::GFP in the genetic background of a standard laboratory strain (BY4742 *MATalpha*) to a panel of wild and industrial genetic backgrounds –e.g. fifty different haploids of the opposite mating type extracted from globally diverse, sequence-validated strains of *Saccharomyces cerivisiae* deposited to the collection of the Royal Netherlands Academy of Arts and Sciences over the past 100 years (CBS; Table 1 and supplement).

**Table 1.**
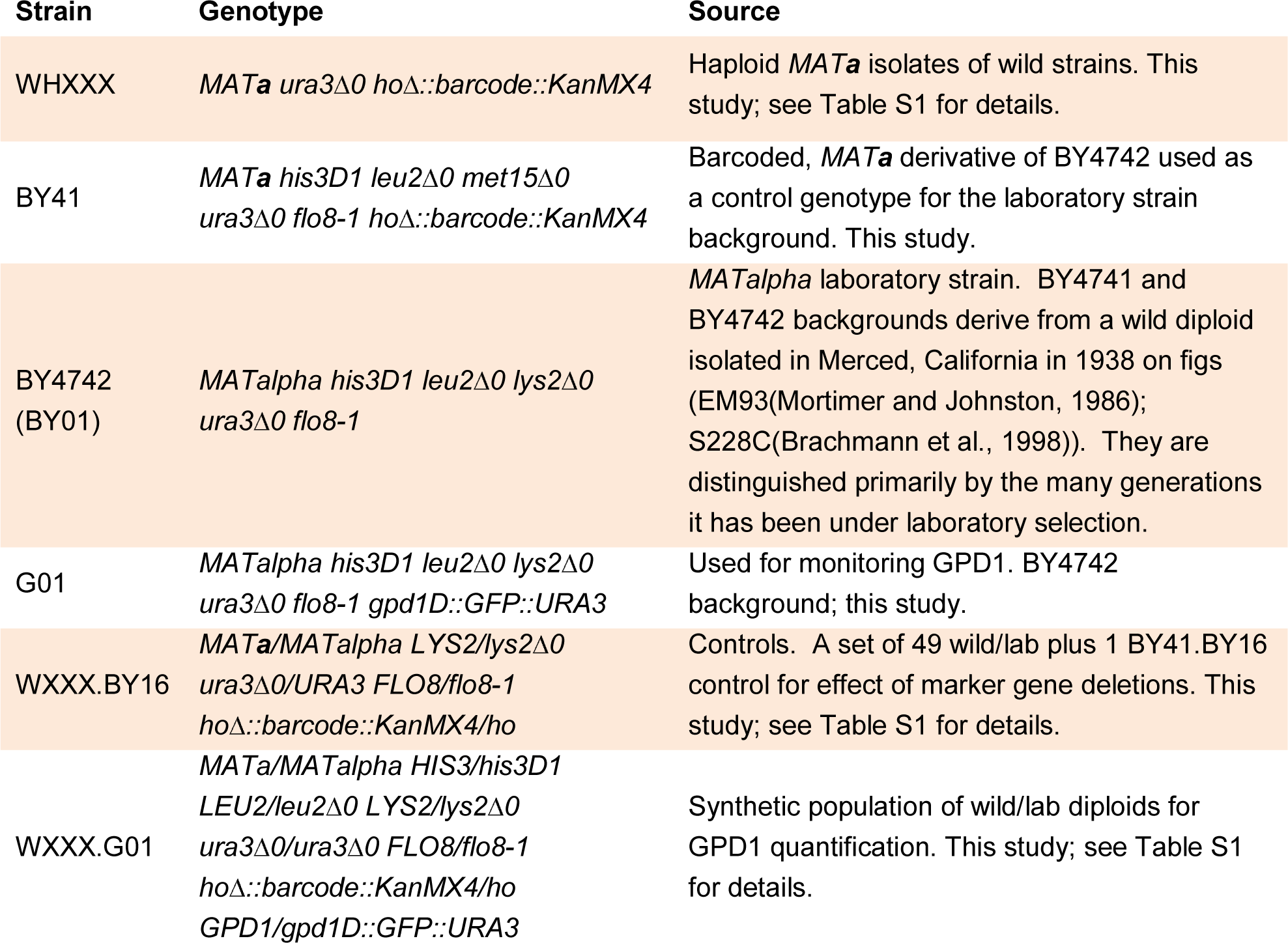
Strains and aliases used in this study. See Table 1–table supplement S1 for details about each of the 49 wild haploid strain derivatives (WHXXX). For brevity, figures are labeled with the wild parent strain number (WXXX; see Table S1 for details).

## Results

### Osmotic stress responses of rapidly dividing cultures

The behavior of single cells before and after their exposure to osmotic stress was followed in several strains by time-lapse video microscopy of monolayer cultures in custom microfluidics devices (Bennett et al., 2008). When cells in exponential growth were exposed to sudden hyperosmotic stress, cell volume decreased, cell division and budding immediately stopped, and daughter cells retracted (Miermont et al., 2013). After a lag period proportional to the severity of the stress GFP fluorescence driven by the *GPD1* promoter began to accumulate in the cytoplasm of surviving cells. Cells that did not accumulate *GPD1*::GFP to high levels did not survive or adapt, developed large vacuoles, and began to die, remaining in view as shrunken cell ghosts. As GFP accumulated to saturation levels in the surviving cells, they adapted to the higher osmotic pressure, resumed cell division, budded and began to divide with a longer half time, producing daughter cells with similarly high fluorescence (Miermont et al., 2013). On the other hand, we measured the viability of each culture and *GPD1*::GFP accumulation per cell using flow cytometry of statistically large numbers of cells from all 50 strains (∼10,000 cells / sample). The rate and extent of mean *GPD1*::GFP accumulation in exponentially growing cultures exposed to hyperosmotic media depended on the severity of the stress and the genetic background of each strain (Figure 1A). Prior to the osmotic stress mean *GPD1*::GFP fluorescence and viability were uncorrelated. After 2 hours in moderate 0.75 M KCl viability decreased and became steeply correlated with accumulated *GPD1*::GFP (Figures 1B and C). As expected, natural variation in the strength of HOG signaling was directly responsible for variation among the strains in osmotic stress survival.

**Figure 1.**
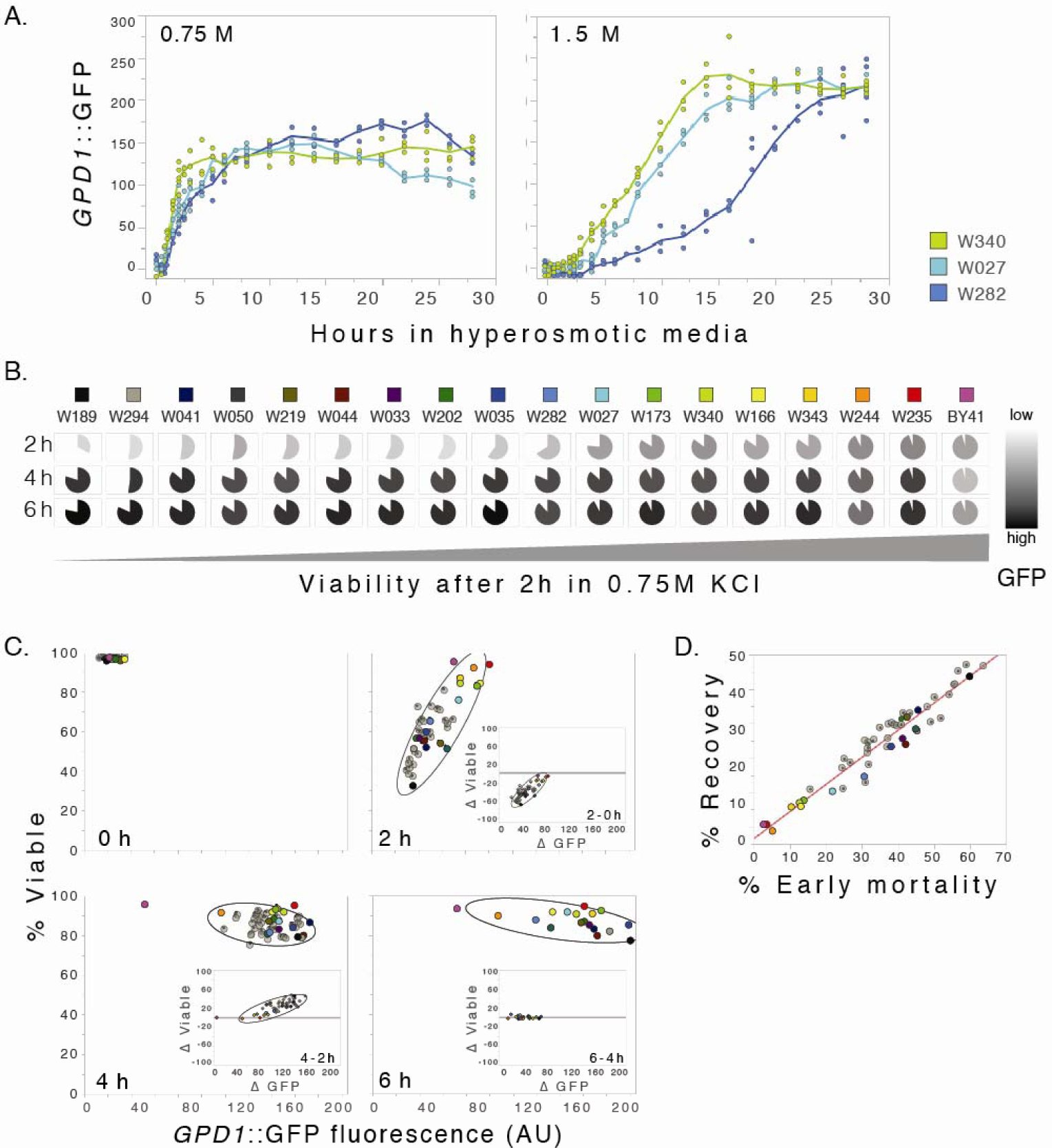
Rate of change in osmotic stress signaling with negative feedback predicts survival and robust recovery of exponential cultures in moderate hyperosmotic stress. A. Time course of mean accumulated *GPD1*::GFP fluorescence in exponential cultures exposed to 0.75 and 1.5 M KCl. Each point represents an independent experimental replicate; curves connect strain means at each time (minimum of 3 replicates per time). In the absence of stress, all strains had high steady-state viability (propidium iodide dye exclusion; range 96.3 – 98.7%; mean 97.6%) and relatively low mean GFP fluorescence indicating low background activity of HOG pathway signaling through the *GPD1* promoter and low *GPD1*::GFP accumulation (range 12.7 – 34.8 AU; mean 18.8 AU). *GPD1*::GFP accumulation reached a steady state by 4 to 6 hours in 0.75 M KCl. B. Pie charts show relative changes in mean viability (shaded area), mortality (white area) and *GPD1*::GFP accumulation (opacity level) of 18 representative strains after 2 hours in 0.75M KCl with strains ordered by increasing viability at 2 hours. Across all 50 strains 2-hour viability was proportional to the 2-hour viability of non-disrupted controls with two intact copies of the *GPD1* gene (*R^2^* = 0.7085; P<0.0001; not shown). C. Relationship between mean *GPD1*::GFP accumulation (AU) and viability in midexponential cultures exposed to 0.75 M KCl for 0, 2, 4, and 6 hours (h). Each data point represents an average of at least three replicates per strain and time (∼10,000 cells/ sample). The ellipses indicate correlations between viability and fluorescence at alpha = 0.95. The inserts show relationships between changes in *GPD1*::GFP and viability over each time interval. D. Negative feedback drove robust recovery of steady-state viability after 4 hours in 0.75 M KCl (robust perfect adaptation; see (Muzzey et al., 2009)). Linear regressions of recovery at 4 hours – recovery = (0.7670) early mortality + 3.49 (*R^2^* = 0.9351; P< 0.0001; 50 strains, shown) and at 6 hours – recovery = (0.7670) early mortality + 3.43 (*R*^2^ = 0.9852; P< 0.0001; 18 strains). Integral feedback control would assure and be assured by perfect adaptation of stress responses, water balance and steady state viability (manuscript in preparation; bioXriv http://dx.doi.org/10.1101/045682). Note that due to the persistence of dead cells in short term cultures (early mortality), 100% recovery of steady-state viability (slope = 1) is not expected over the course of the experiment. The data are fit well by a model whereby dead cells remain and surviving cells in all strains undergo 3 cell divisions (not shown).

### Negative feedback drives a robust recovery

The initially strong positive correlation between variation in *GPD1*::GFP accumulation and variation in viability reversed as cells adapted and began to divide (Figure 1C; 4 hours). This distinguished an early phase (0 – 2 hours) of the response when viability decreased markedly and acute HOG signaling promoted osmotic stress survival and a later phase (2 – 4 hours) when viability recovered but became negatively correlated with HOG signaling and *GPD1*::GFP accumulation. The switch from positive to negative correlations might have indicated that stronger HOG signaling, initially beneficial, suddenly caused lower viability. However, negative feedback controls, occurring at many levels and timescales, are present in essentially all of the varied mechanisms that act in concert to increase intracellular glycerol and restore water balance. We think it more likely that negative feedback increased signaling in the surviving cells of the less viable strains. For example (1) unequal water pressures activate osmotic stress sensors, glycerol channels and other pressure-sensitive components whose activities control and depend on water balance (e.g. see Figure 5 in Hohmann 2002 (Hohmann, 2002; Saito and Posas, 2012)), (2) GPD1 indirectly controls and is controlled by osmotic stress-sensitive kinases that respond to upward and downward changes in water balance (Lee et al., 2012), and (3) nuclear Hog-1 MAP kinase increases the transcription of phosphatases that restore its own cytoplasmic localization and basal activity (Jacoby et al., 1997; Muzzey et al., 2009; Wurgler-Murphy et al., 1997).

Consistent with acting negative regulation, there was a strong and highly significant correlation between early mortality (0 – 2 hour decreases in viability) and later accumulations of *GPD1*::GFP (2 – 4 hours; Table 2). We reasoned that cells and strains that adapt quickly experience lower and less sustained effects of osmotic stress (e.g. water loss) with more rapidly attenuated HOG pathway activity and lower *GPD1*::GFP accumulation. Conversely, surviving cells of strains that were slower to adapt and less viable would experience higher and more sustained osmotic (and likely other) stress(es). Prolonged osmotic stress would maintain HOG signaling and *GPD1* transcription – which is also activated by general stress responses(Boy-Marcotte et al., 1998) – further promoting *GPD1*::GFP accumulation (e.g. a negative feedback regulation of viability by general stress responses). Indeed, even as *GPD1*::GFP and viability became negatively correlated, their *rates of change* remained positively correlated (Figure 1C, 2 – 4 hours and insets; Table 2), prompting a parsimonious interpretation that osmotic stress signaling promotes adaptation and viability during both the initial and recovery phases of the response.

**Table 2.** Negative feedback between rates of change in mean *GPD1*::GFP accumulation and viability among strains. Correlations confirm causality between rates of change in *GPD1*::GFP accumulation and viability within (upper 3 rows) and between 2 hour time intervals (below). Changes occurring in earlier intervals are listed first. To control for potential deviations from normality, both parametric (Pearson’s) and non-parametric (Spearman’s) pairwise correlations are shown. As in Figure 1 all 50 strains were tested at 0, 2, and 4 hours and 18 strains were tested at 6 hours (mean values represent a minimum of 3 replicates per strain). Significant comparisons are in bold (JMP statistical software, SAS Institute; Cary, NC).

| N | Variable | Interval (hrs) | Variable | Interval (hrs) | Pearson's r | Spearman's r | Prob>\|r\| |  |
| --- | --- | --- | --- | --- | --- | --- | --- | --- |
|  |  |  |  |  |  |  | Pearson's | Spearman's |
| 50 | $\Delta GPD1::GFP$ | 0 – 2 | Δviability | 0 – 2 | 0.8235 | 0.7725 | **<.0001** | **<.0001** |
| 50 | $\Delta GPD1::GFP$ | 2 – 4 | Δviability | 2 – 4 | 0.7739 | 0.7217 | **<.0001** | **<.0001** |
| 18 | $\Delta GPD1::GFP$ | 4 – 6 | Δviability | 4 – 6 | -0.2354 | -0.1992 | 0.3470 | 0.4282 |
| 50 | Δviability | 0 – 2 | $\Delta GPD1::GFP$ | 2 – 4 | -0.7867 | -0.7411 | **<.0001** | **<.0001** |
| 50 | Δviability | 0 – 2 | Δviability | 2 – 4 | -0.9670 | -0.9503 | **<.0001** | **<.0001** |
| 50 | $\Delta GPD1::GFP$ | 0 – 2 | $\Delta GPD1::GFP$ | 2 – 4 | -0.7685 | -0.7697 | **<.0001** | **<.0001** |
| 50 | $\Delta GPD1::GFP$ | 0 – 2 | Δviability | 2 – 4 | -0.8082 | -0.7696 | **<.0001** | **<.0001** |
| 18 | Δviability | 0 – 2 | Δviability | 4 – 6 | -0.1407 | -0.2178 | 0.5777 | 0.3854 |
| 18 | Δviability | 0 – 2 | $\Delta GPD1::GFP$ | 4 – 6 | -0.3704 | -0.2549 | 0.1303 | 0.3073 |
| 18 | $\Delta GPD1::GFP$ | 0 – 2 | Δviability | 4 – 6 | -0.0456 | -0.0464 | 0.8573 | 0.8548 |
| 18 | $\Delta GPD1::GFP$ | 0 – 2 | $\Delta GPD1::GFP$ | 4 – 6 | -0.4319 | -0.4572 | 0.0735 | 0.0565 |
| 18 | Δviability | 2 – 4 | Δviability | 4 – 6 | -0.0100 | 0.0733 | 0.9685 | 0.7726 |
| 18 | Δviability | 2 – 4 | $\Delta GPD1::GFP$ | 4 – 6 | 0.4316 | 0.3333 | 0.0737 | 0.1765 |
| 18 | $\Delta GPD1::GFP$ | 2 – 4 | Δviability | 4 – 6 | 0.1400 | 0.1207 | 0.5796 | 0.6332 |
| 18 | $\Delta GPD1::GFP$ | 2 – 4 | $\Delta GPD1::GFP$ | 4 – 6 | 0.3731 | 0.3602 | 0.1273 | 0.1421 |

By 4 hours all strains had adapted to a new steady state in 0.75M KCl and later viability remained largely unchanged (Figure 1C inset, lower right). Interestingly, initial decreases in steady-state viability (0 – 2 hour mortality) were almost perfectly restored by 4 hours (Figure 1D) and, remarkably, by 6 hours early mortality and recovery were over 98% correlated (*^R^2* = 0.9852, P<0.0001, N=18; see legend Figure 1D). The biological robustness of adaptation and complete recovery of steady state viability further support the idea that negative feedback restores viability through continued activation of stress responses. Indeed, the continued accumulation of *GPD1* and glycerol – directly responsible for restoration of water balance and reduction of osmotic stress –suggests that intracellular glycerol concentrations integrate the cumulative activities of many facets of the osmotic stress response (e.g. provides a plausible biological mechanism for “integral feedback” that virtually assures perfect adaptation(Muzzey et al., 2009; Yi et al., 2000); manuscript in review). However, despite their resilience, strains that were relatively slower to adapt would be ultimately less fit than rapidly adapting strains due to their higher death rate, slower recovery, and lower viabilities before and after adaptation.

### Extreme stress resistance of older cultures

By contrast with exponential cultures, when the aging yeast cultures (post-diauxic) were exposed to hyperosmotic media they survived and adapted after long periods in unprecedented conditions (Movies 1 and 2). As aging cultures deplete available glucose in their media they undergo a metabolic change called the diauxic shift (Galdieri et al., 2010). During post-diauxic growth stress response proteins accumulate, cell division slows and then stops, and cells enter quiescence (Gray et al., 2004). Remarkably, post-diauxic cultures survived up to 5 weeks in 3 M KCl (41/50 strains). They could not adapt and did not grow in 3 M KCl, but recovered rapidly and grew when plated on fresh isotonic media (Figure 2B). When we tested their limits of adaptation in increasing concentrations of KCl all but one strain could grow on 2.6 M KCl media and three strains could grow on media containing 2.9 M KCl (Table 3). We are unaware of previous reports of such extreme osmotic stress survival or adaption limits for budding yeast of any growth stage or genotype.

**Table 3.** Growth of post-diauxic cells at unprecedented limits of adaptation. Shown are concentrations of agar media on which post-diauxic strains could grow and form colonies.

| [KCl] M | Wild/lab (*GPD1*) diploids* |
| --- | --- |
| 2.0 | W455 |
| 2.6 | W027, W035, W167, W202, W203, W242, W285, W454 |
| 2.7 | W033, W041, W042, W134, W136, W150, W166, W178, W195, W215, W217, W219, W235, W248, W282, W291, W292, W294, BY41 |
| 2.8 | W037, W044, W050, W153, W157, W163, W164, W179, W189, W206, W238, W244, W245, W249, W255, W276, W301, W340 |
| 2.9 | W173, W211, W343 |

**Figure 2.**
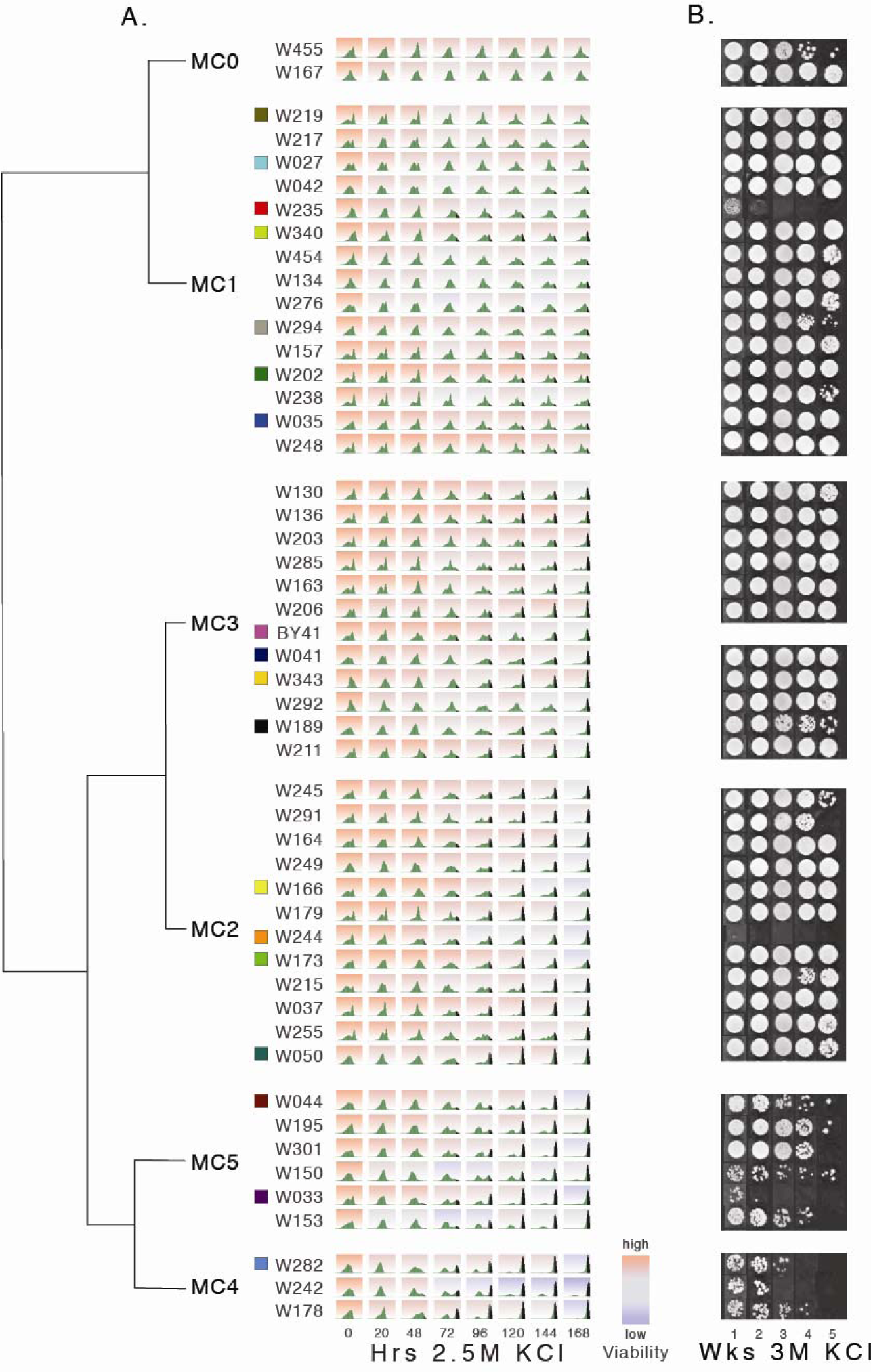
Continuous variation in survival and rates of accumulation of *GPD1*::GFP fluorescence of postdiauxic cultures during severe hyperosmotic osmotic stress. A. Strains were clustered (MC0 – MC5) and ranked according to their rates of accumulation of *GPD1*::GFP fluorescence (see Table 5 and methods). A representative distribution of *GPD1*::GFP accumulation (green) and relative survival red (99.7% viability) to blue (11.7% viability) is given for each strain and time point (4-15 fold replication). Distributions of cells above the 89^th^ percentile (top 11%) are shown in black. Ranking was sequential from 1 (top) to 50 (bottom). Prior to osmotic challenge (0 hours in 2.5 M KCl) steady-state viabilities were uniformly high (range 93.0 – 99.6%; mean 98.2%). Strains are color-coded as in Figure 1C for comparison of exponential and post-diauxic cultures. B. Relative viability of post-diauxic cultures (WXXX.BY01 controls) incubated in 3 M KCl before plating on iso-osmolar media. Platings were re-ordered according to the ranked signaling behavior given in Figure 2A.

### Heterogeneity of cells in older cultures

By contrast with cultures in exponential growth, in post-diauxic growth the genetically identical cells within each strain and culture were surprisingly heterogeneous in their size, shape and signaling behaviors (Figure 2). Neither total *GPD1*::GFP fluorescence nor rates of change in fluorescence was strongly correlated with viability. After several hours in 2.5 M KCl, *GPD1*::GFP increased sharply in one group of cells as they began to divide. More surprising, another group of cells induced *GPD1*::GFP to high levels, started to divide and then popped, killing both the mother and daughter (Movie 3). Other cells had slower signaling and cell division while the most ‘cautious’ groups of cells failed to signal or divide but remained in a cellular state of static viability without dividing.

The unique signaling trajectories of most strains were highly reproducible. We used machine learning to assign the different behaviors of the cells in each sample to four Gaussian distributions (G_0_-G_3_) described by eight parameters – means and covariances – numbered according to their increasing levels of fluorescence (Figure 2–figure supplement 1). Only the mean level of *GPD1*::GFP pre-accumulated into cells of the G_3_ distribution of each strain during post-diauxic growth – *prior* to the osmotic challenge and therefore unrelated to osmotic stress signaling – predicted survival at any time. The amount of *GPD1*::GFP in G_3_ cells at time 0 predicted early but not later viability and this relationship was better fit by 2^nd^ order quadratic rather than linear functions of *GPD1*::GFP (Table 4), showing that early survival was higher in strains with intermediate G_3_ accumulations (more variation explained and lower mean square errors). Despite the fine-scaled characterization of osmotic stress signaling behaviors of the different groups of cells in each strain, none of the distributions learned by the Gaussian mixture model, neither pre-accumulated G_3_, total *GPD1*::GFP fluorescence, nor stress-induced *GPD1*::GFP in any distribution, embodied features of osmotic stress signaling important for later survival.

**Table 4.** Osmotic stress signaling behavior (rank) predicted early and late viability of post-diauxic cultures in osmotic stress. Least squares predictions of early and late viability by linear and 2^nd^ order quadratic fits of fluorescence pre-accumulated into the G3 Gaussian at time 0 (G3_0) and ranked signaling behavior of 50 strains. The Bonferroni cutoff at the 0.05 level, based on 4 tests per data set, was 0.0125 (JMP statistical software, SAS Institute; Cary, NC). Significant fits with lowest root mean squared errors and highest fraction of variation explained (*R*^2^) shown in bold, predicted values for optimum (x) and value at optimum (y) for non-significant fits are shown for comparison.

|  | G<sub>3</sub> fluorescence (AU)<br>at time 0 |  | Signaling (rank) |  |
| --- | --- | --- | --- | --- |
|  | quadratic | linear | quadratic | linear |
| **0 hours, 0M KCl** |  |  |  |  |
| probability > F | **0.0006** | 0.0016 | 0.9629 | 0.8454 |
| R_square | **0.2715** | 0.1891 | 0.0016 | 0.0008 |
| root_mean_square_error | **1.3935** | 1.4548 | 1.6313 | 1.6149 |
| max_viability at optimum (%) | **98.8** |  | NS 98.2 |  |
| optimum AU or rank | **2626.6** |  | NS 31.9 |  |
| **20 hours, 2.5M KCl** |  |  |  |  |
| probability > F | **< 0.0001** | < 0.0001 | **0.0055** | 0.6023 |
| R_square | **0.4286** | 0.3138 | **0.1987** | 0.0057 |
| root_mean_square_error | **7.0949** | 7.6940 | **8.4022** | 9.2615 |
| max_viability at optimum (%) | **86.7** |  | **86.3** |  |
| optimum AU or rank | **2662.5** |  | **24.4** |  |
| **48 hours, 2.5M KCl** |  |  |  |  |
| probability > F | **< 0.0001** | 0.0003 | **0.0013** | 0.9159 |
| R_square | **0.3148** | 0.2371 | **0.2464** | 0.0002 |
| root_mean_square_error | **7.6281** | 8.1323 | **8.1679** | 9.3096 |
| max_viability at optimum (%) | **82.2** |  | **82.9** |  |
| optimum AU or rank | **2622.3** |  | **25.7** |  |
| **72 hours, 2.5M KCl** |  |  |  |  |
| probability > F | **0.0010** | 0.0033 | **0.0062** | 0.8464 |
| R_square | **0.2556** | 0.1660 | **0.1943** | 0.0008 |
| root_mean_square_error | **9.9877** | 10.4608 | **10.3903** | 11.4501 |
| max_viability at optimum (%) | **73.5** |  | **74.3** |  |
| optimum AU or rank | **2586.8** |  | **25.1** |  |
| **96 hours, 2.5M KCl** |  |  |  |  |
| probability > F | **0.0060** | 0.0018 | **0.0065** | 0.0954 |
| R_square | **0.1956** | 0.1862 | **0.1927** | 0.0569 |
| root_mean_square_error | **8.4186** | 8.3789 | **8.4338** | 9.0201 |
| max_viability at optimum (%) | **71.7** |  | **71.0** |  |
| optimum AU or rank | **3389.8** |  | **21.3** |  |

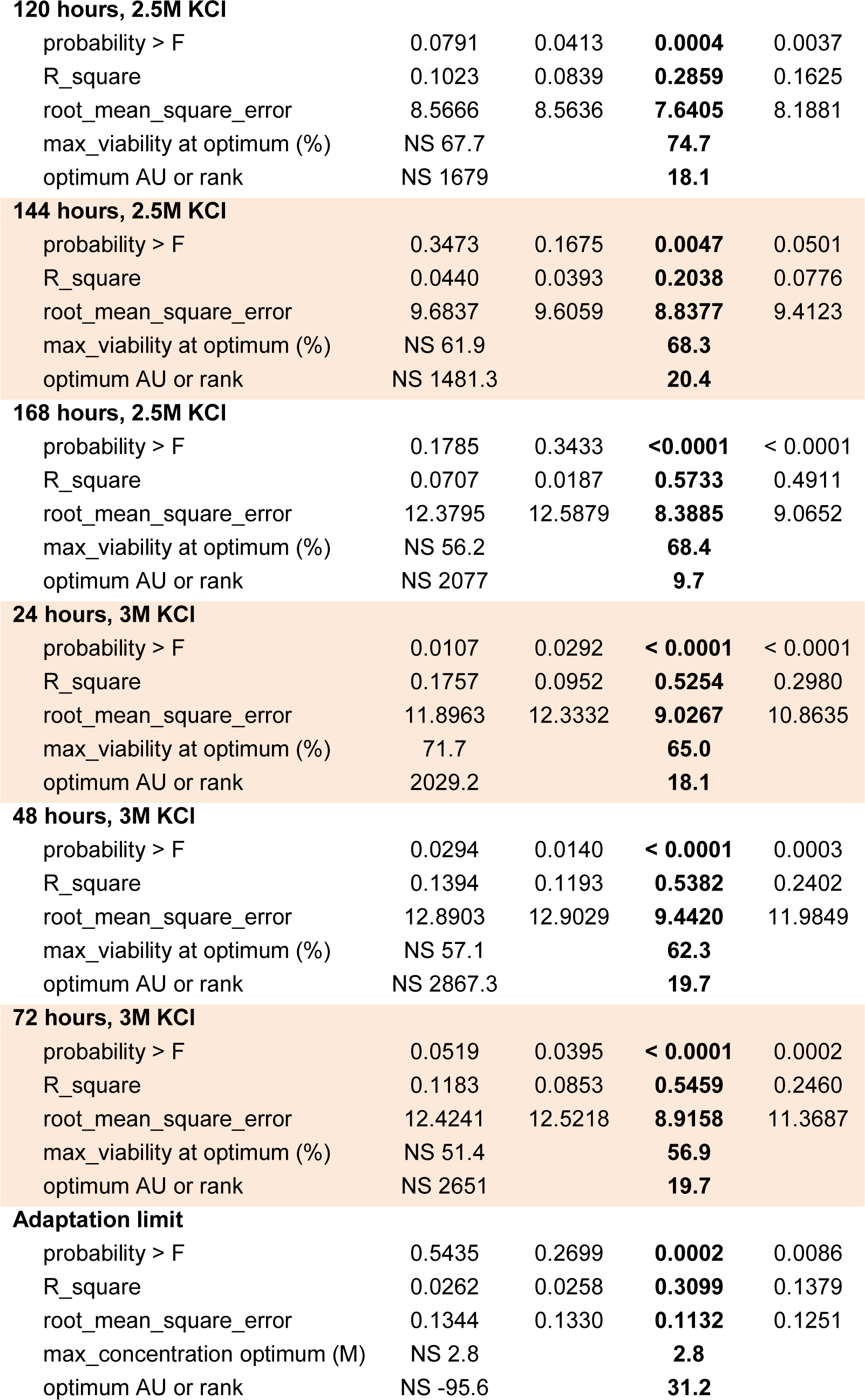

**120 hours, 2.5M KCI**
|  |  |  |  |  |
| --- | --- | --- | --- | --- |
| probability > F | 0.0791 | 0.0413 | **0.0004** | 0.0037 |
| R_square | 0.1023 | 0.0839 | **0.2859** | 0.1625 |
| root_mean_square_error | 8.5666 | 8.5636 | **7.6405** | 8.1881 |
| max_viability at optimum (%) | NS 67.7 |  | **74.7** |  |
| optimum AU or rank | NS 1679 |  | **18.1** |  |
| **144 hours, 2.5M KCI** |  |  |  |  |
| probability > F | 0.3473 | 0.1675 | **0.0047** | 0.0501 |
| R_square | 0.0440 | 0.0393 | **0.2038** | 0.0776 |
| root_mean_square_error | 9.6837 | 9.6059 | **8.8377** | 9.4123 |
| max_viability at optimum (%) | NS 61.9 |  | **68.3** |  |
| optimum AU or rank | NS 1481.3 |  | **20.4** |  |
| **168 hours, 2.5M KCI** |  |  |  |  |
| probability > F | 0.1785 | 0.3433 | **<0.0001** | < 0.0001 |
| R_square | 0.0707 | 0.0187 | **0.5733** | 0.4911 |
| root_mean_square_error | 12.3795 | 12.5879 | **8.3885** | 9.0652 |
| max_viability at optimum (%) | NS 56.2 |  | **68.4** |  |
| optimum AU or rank | NS 2077 |  | **9.7** |  |
| **24 hours, 3M KCI** |  |  |  |  |
| probability > F | 0.0107 | 0.0292 | **< 0.0001** | < 0.0001 |
| R_square | 0.1757 | 0.0952 | **0.5254** | 0.2980 |
| root_mean_square_error | 11.8963 | 12.3332 | **9.0267** | 10.8635 |
| max_viability at optimum (%) | 71.7 |  | **65.0** |  |
| optimum AU or rank | 2029.2 |  | **18.1** |  |
| **48 hours, 3M KCI** |  |  |  |  |
| probability > F | 0.0294 | 0.0140 | **< 0.0001** | 0.0003 |
| R_square | 0.1394 | 0.1193 | **0.5382** | 0.2402 |
| root_mean_square_error | 12.8903 | 12.9029 | **9.4420** | 11.9849 |
| max_viability at optimum (%) | NS 57.1 |  | **62.3** |  |
| optimum AU or rank | NS 2867.3 |  | **19.7** |  |
| **72 hours, 3M KCI** |  |  |  |  |
| probability > F | 0.0519 | 0.0395 | **< 0.0001** | 0.0002 |
| R_square | 0.1183 | 0.0853 | **0.5459** | 0.2460 |
| root_mean_square_error | 12.4241 | 12.5218 | **8.9158** | 11.3687 |
| max_viability at optimum (%) | NS 51.4 |  | **56.9** |  |
| optimum AU or rank | NS 2651 |  | **19.7** |  |
| **Adaptation limit** |  |  |  |  |
| probability > F | 0.5435 | 0.2699 | **0.0002** | 0.0086 |
| R_square | 0.0262 | 0.0258 | **0.3099** | 0.1379 |
| root_mean_square_error | 0.1344 | 0.1330 | **0.1132** | 0.1251 |
| max_concentration optimum (M) | NS 2.8 |  | **2.8** |  |
| optimum AU or rank | NS -95.6 |  | **31.2** |  |

### Continuous variation in stress responses

In order to map osmotic stress-responsive signaling onto survival more directly, we clustered the osmotic stress signaling trajectories of each strain using state vectors to describe directly *GPD1*::GFP distributions of cells in each culture unbiased by Gaussian assumptions or approximations. Based on their shared and strain-specific (heritable) signaling behaviors, the 50 strains rapidly converged onto two large groups made up of six mean clusters (Figure 2). Each strain was further ordered within and between mean clusters based on their clustering statistics (Table 5), with their rank order describing increasingly rapid accumulations of *GPD1*::GFP and ‘reckless’ signaling (Figure 2-figure supplement 2). Tellingly, both mean cluster (Figure 3B) and rank (Figure 3C) predicted viability over time (Table 4), thereby confirming the biological relevance of ‘cautious’ versus ‘reckless’ osmotic stress signaling, validating our clustering method and supporting the role of natural osmotic stress signaling differences between strains in shaping variation in fitness during osmotic stress.

**Table 5.**
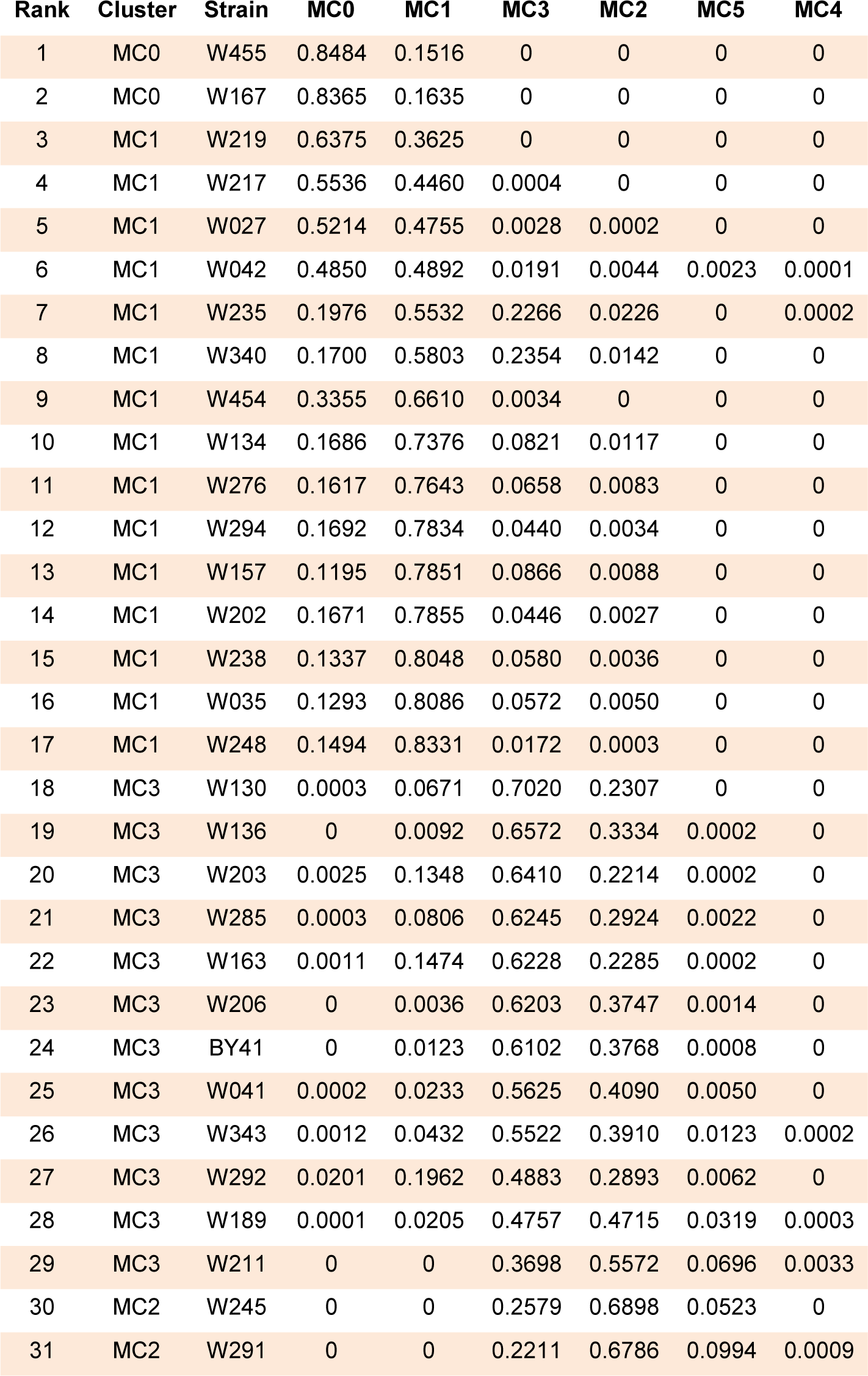
Clustering statistics used to rank signaling behavior. Statistics showing the fraction of 17,000 permutations in which strains were clustered with at least 50% of the other strains in each mean cluster. These data were used to rank total signaling behaviors from most cautious (1) to most reckless (50) based on the fraction of time each strain was associated with its mean cluster (characteristic of that cluster). See Figure 2.

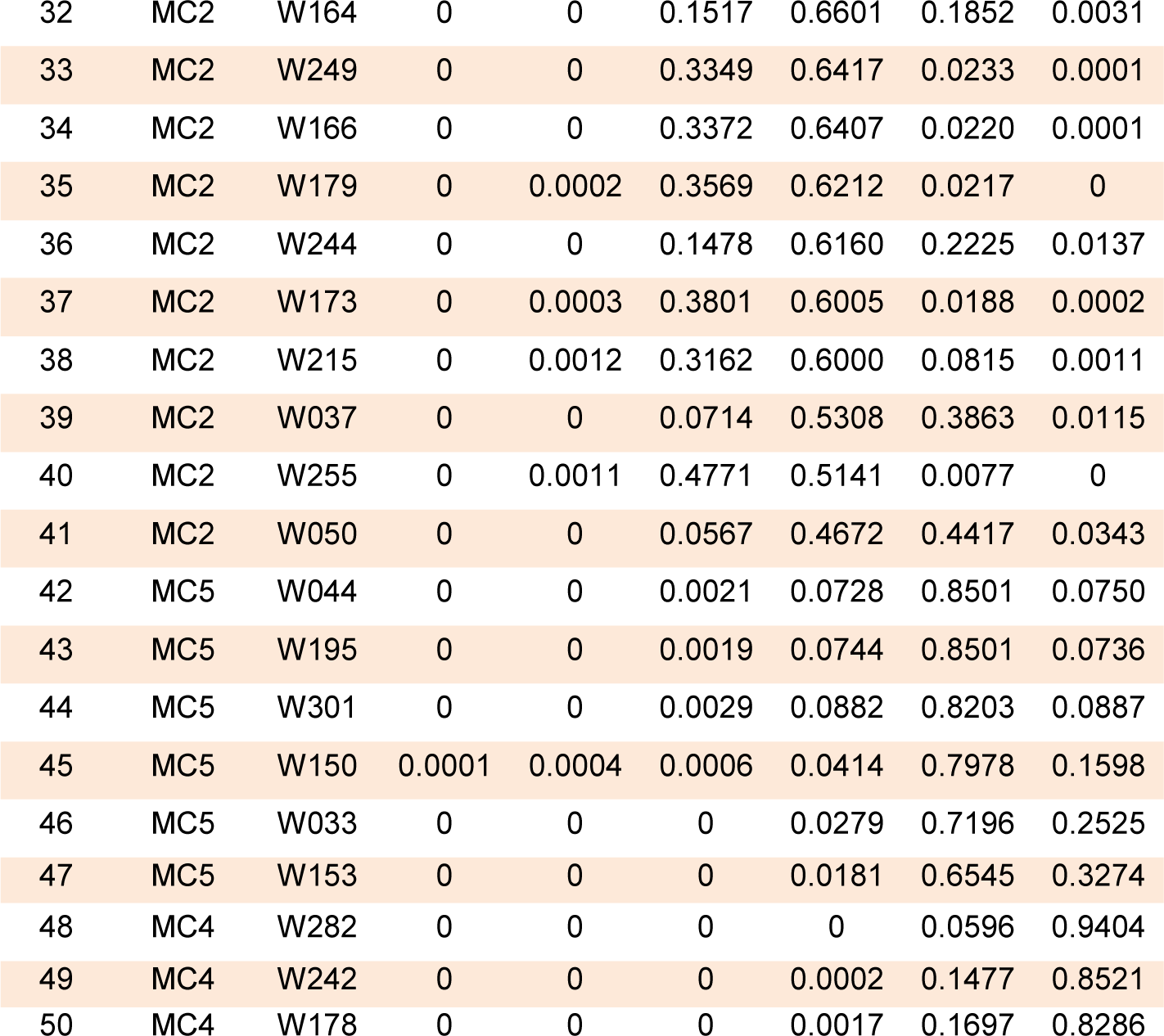

**Figure 3.**
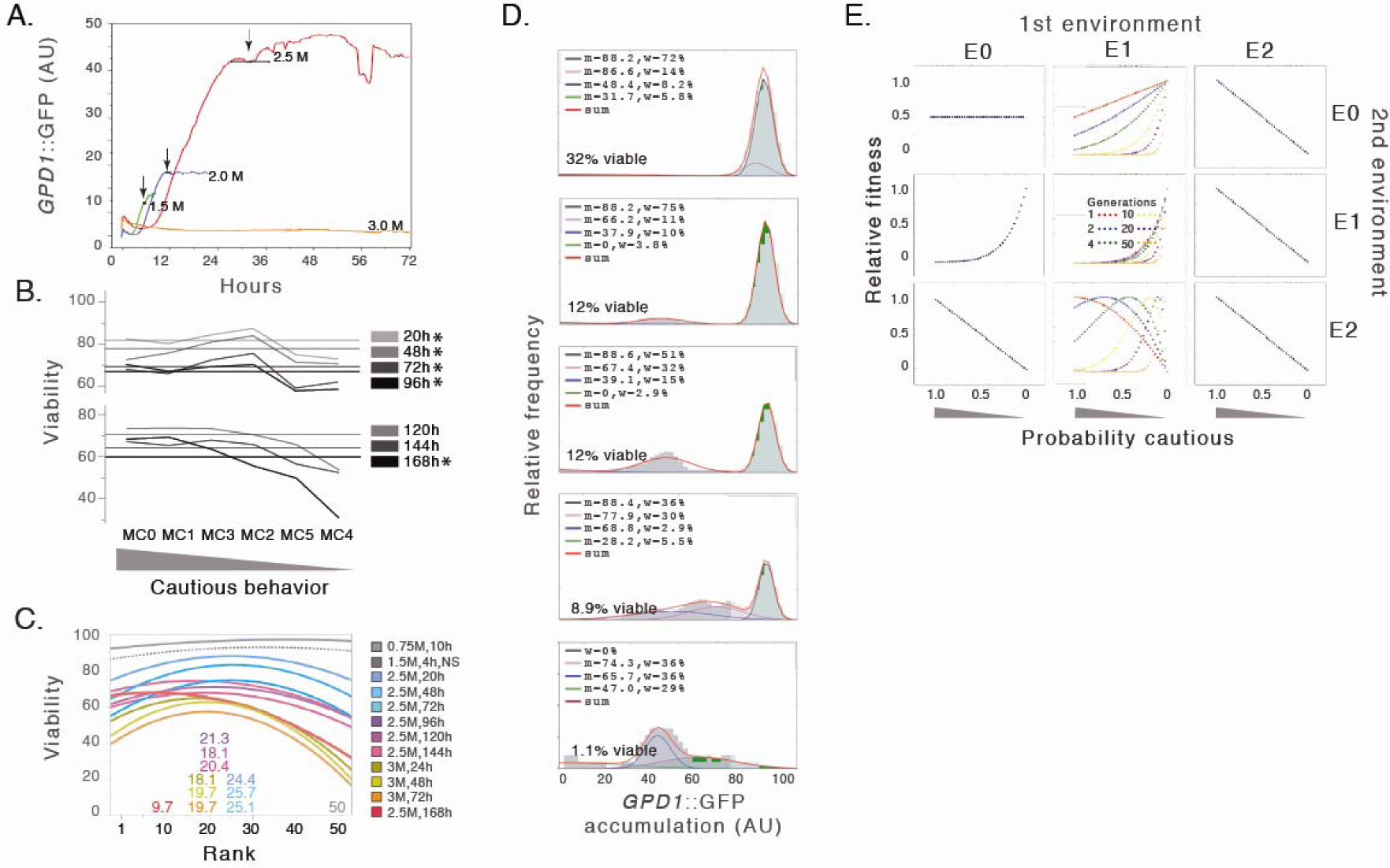
Modeling a heritable probability of cautious behavior (bet hedgers) produces observed variation in relative fitness and survival. A. The most cautious behavior (static viability) of post-diauxic cells from strain W027 exposed to 3 M in microfluidic chambers. Individual cell behaviors mirror population behaviors measured by flow cytometry – e.g longer lag periods and increased accumulations of *GPD1*::GFP with increasing osmotic stress. Colored traces indicate accumulated fluorescence (AU X 10^−2^) in representative cells at 1.5 (green), 2.0 (blue), 2.5 (red) and 3.0 M KCl (yellow). Arrows indicate average time to the first cell division +/- standard deviations. B. Mean cluster predicts viability in different osmotic environments. Average viabilities among mean clusters for post-diauxic cultures in 2.5 M KCl for indicated times. Horizontal lines show overall average viability at each time point across all 50 strains. The numbers of strains in each mean cluster are 2 (MC0), 15 (MC1), 12 (MC2), 12 (MC3), 3 (MC4), 6 (MC5; see Figure 2). Asterisks indicate significance at the ≤0.05 level by ANOVA or, where appropriate because variances were unequal, Welch’s ANOVA (JMP statistical software, SAS Institute, Cary, NC). C. Rank predicts viability in different osmotic environments. Shown are the best-fit curves for viability by rank in each of the environments listed (2^nd^ order quadratic relationships; Table 4). The optimal signaling strategy (rank) shifted from higher (50; most reckless) to lower (most cautious) as the environment became progressively more severe. Dashed lines indicate the relationship between rank and viability was only marginally significant in that environment. D. Cells with the most aggressive signaling began to die after long periods in severe stress leaving increasing fraction of cells with lower *GPD1*::GPF accumulations. Shown are distributions of accumulated *GPD1*::GFP fluorescence (AU) and viability in replicate cultures of W242 (rank 49) in 5 replcate cultures after 168 hours in 2.5 M KCl. Mean (x), standard deviation (std), and weight (w; the fraction of cells in each distribution) are given. Sum (red) shows the cumulative fit of the 4 learned Gaussians. E. A simple bet hedging model with heritable proportions of cautious and reckless cells produces observed variation in survival. Bet hedging strategy P was defined as the probability of cautious cells for 0 ≤ P ≤ 1. Relative fitness was measured for all strategies after 10 generations in each environment. All nine possible 2-state environmental shifts between three general osmotic stress environments were considered: permissive (E0; all cells grow equally well), restrictive (E1; reckless cells divide, cautious cells survive without dividing), and killing (E2; reckless cells die, cautious cells survive without cell division). Intermediate strategies (0 < P < 1; i.e. bet hedging) were most fit only when the environment shifted from moderate to more severe (E1 -> E2). When E1 was the first environment, the optimum strategy P depended on generation number.

### Evidence for bet hedging

As cautious and reckless behaviors were found both within and between strains, we wondered whether bet hedging, the expression of alternate, conditionally-adaptive phenotypes within a clone of genetically identical organisms, could explain the observed variation in osmotic stress signaling and survival (King and Masel, 2007; Meyers and Bull, 2002; Philippi and Seger, 1989; Ratcliff et al., 2014; Simons, 2011). By contrast with previously described stochastic bet hedging, where pre-adapted stress-resistant cells occur at generally low frequencies in clonal populations (De Jong et al., 2011; Levy et al., 2012), ranked signaling responses induced by osmotic stress were uncorrelated with the pre-adapted (G_3_) resistance acquired during post-diauxic growth and already present at time 0. Strain viabilities varied depending on their rank and the severity of the osmotic environment (Table 4). Milder conditions favored higher-ranked strains with more aggressive osmotic stress signaling strategies, but with increasing time and harsher conditions more cautious strains and behaviors became more fit. For example, W178 at rank 50 was most viable in moderate 0.75 M KCl, but the optimum shifted to rank 25 after 20 hours in 2.5 M KCl, 20 after 72 hours, and 9.6 after 168 hours (1 week). After 168 hours, viability had decreased most among the most reckless, highest-ranked strains (Figures 3B and C). To confirm that the increasing survival of cautious strains was correlated with their experience of osmotic stress and not simply time in culture, we again incubated cultures in 2.5 M KCl for 168 hours, but they were first exposed to a mild pre-stress (2 hours in 0.5 M KCl) to pre-induce osmotic stress proteins. If optimum rank depended solely on time in high KCl, e.g. independent of the degree of stress “experienced” by the different strains, then it should be unaffected by the short pre-stress. However, optimum rank shifted toward more reckless behaviors (rank 9.6 to rank 18; P<0.0001) and viability increased by ∼10% in response to the pre-stress. This suggests pre-treated cells experienced lower osmotic stress and that the stress itself directly determined the optimum signaling behavior. The strong correlation between relative fitness and signaling behavior with increasing osmotic stress (see below) provides strong empirical evidence for bet hedging (Simons, 2011).

After 168 hours in 2.5 M KCl, the most reckless cells in the highest-ranking strains began to selectively die and disappear. For example, among replicate cultures of strain W242 (rank 49) those with lower viability had fewer cells with high accumulations of *GPD1*::GFP, smaller G_3_ distributions, and correspondingly larger distributions of cells with lower mean *GPD1*::GFP (Figure 3D). A selective loss of cells with the highest accumulations of *GPD1*::GFP could indicate that *GPD1*::GFP levels simply decrease over time. However, G_3_ distributions were stable over most time points and in most strains (e.g. Figure 2-figure supplement 1). We think it more likely that after 168 hours the most aggressive cells attempted to divide and popped (as in Movie 3), preferentially decreasing the G_3_ distribution relative to the other distributions. Rapid signaling, adaptation and recovery of cell division, a fitness advantage in mild conditions, becomes a liability in severe or prolonged osmotic stress. On the other hand, static viability would dramatically reduce evolutionary fitness in normal environments but would allow more cautious cells and strains to survive extreme hyperosmotic stress.

### Evolution of bet hedging

As cautious and reckless strains reliably express a range of cells with different behaviors and fitness depending on the environment, we wondered whether a simple, 2-state bet hedging model including heritable proportions of cautious and reckless cell types could account for the observed variation in osmotic stress signaling and explain the complex relationship between rank and viability. Assuming aggressive osmotic stress signaling with rapid recovery and resumption of growth is the default, ancestral behavior, we hypothesized a heritable probability of cautious signaling and behavior arose in response to the unpredictable severity and duration of potentially lethal osmotic environments. To test this idea, we modeled the relative fitness of strains with different signaling strategies after several generations of growth, including abrupt changes between three osmotic stress environments that discriminate cautious versus reckless behavior. These were: (E0) a permissive environment in which both cautious and reckless cells grow equally well, (E1) a restrictive environment approximating moderate osmotic stress where reckless cells divide and cautious cells survive without dividing, and (E2) a killing osmotic stress where reckless cells die and cautious cells survive without cell division.

We modeled a heritable probability of daughters with cautious signaling and behavior and asked whether it could evolve (P: 0 ≤ P ≤ 1). Our model calculates the relative fitness (cell numbers) of different strategies after several generations under each of the 9 possible environmental shifts between the three different selective environments (Figure 3E). Most combinations of environments favor an optimum strategy of either all cautious (P = 1) or all reckless cell types (P = 0). Strictly intermediate strategies and bet hedging (P: 0 < P < 1) prevailed only when the osmotic environment changed from moderate to more severe (E1 -> E2) with the optimum strategy P depending on the number of generations in the first environment. Shorter lag periods – corresponding to less severe osmotic conditions – and more cell divisions in E1 initially favor lower P and a higher proportion of reckless cells. Longer lag periods – corresponding to more severe conditions and fewer cell divisions – favor higher P and more cautious cells. Such worsening conditions are common in nature (for example, during fermentation or on drying fruit). Indeed, as predicted by the model, we observed that lower-ranked strains with more cautious signaling behaviors, longer lag periods and fewer attempted cell divisions were increasingly fit over time in an increasing severity of osmotic stress. This simple model provides a conceptual framework for understanding how a heritable frequency of bet hedgers can be tuned by evolution in different patterns of environmental stress.

## Discussion

In order to understand the evolutionary trajectories of populations and species we need to understand the effects of natural genetic variation on mechanisms of development and expression of phenotypic variation. Mapping genetic variation to the spectrum of attributes and behaviors upon which selection acts defines population-level properties such as evolvability (the capacity to evolve), robustness or canalization (the capacity to withstand genetic and environmental perturbation), and norms of reaction (optimization, within a given genotype, of phenotypic responses to different environments) (Kirschner and Gerhart, 1998; Meyers and Bull, 2002; Rutherford, 2000; West-Eberhard, 2003). Predictive plasticity, the ability to sense and respond appropriately to environmental challenge is a hallmark of environmental stress responses. On the other hand, diversified bet hedging, the stochastic expression of phenotypic diversity, arises when environmental signals preceding lethal stresses are unreliable or sudden. Recent theoretical papers discuss ecological forces that favor the evolution of predictive plasticity and diversified bet hedging strategies (Arnoldini et al., 2012; Donaldson-Matasci et al., 2013).

Bet hedging in microorganisms was originally thought to arise almost exclusively through stochastic switching of a small fraction of cells independent of environmental cues (Levy et al., 2012; Ratcliff et al., 2014). However a recent survey of morphological variation among genetically identical cells from 34 different yeast strains demonstrates that differences between strains and traits in phenotypic noise is genetically encoded and if adaptive, could therefore evolve (Yvert et al., 2013). Here we report that even when yeast cells can sense and respond appropriately to unexpected episodes of environmental stress – a classic form of predictive plasticity – to safely resume cell division they must also anticipate the imperfectly known trajectory and duration of environmental change. The yeast osmotic stress response thus combines predictive plasticity with generation of phenotypic diversity and bet hedging to optimize how aggressively and over what time course osmotic stress responses unfold. Indeed, the theoretical models predict the entanglement of plastic developmental responses with bet hedging and circumscribe the ecological settings under which they would evolve (Arnoldini et al., 2012; Donaldson-Matasci et al., 2013). The first known example of combined predictive plasticity and bet hedging in a microorganism is the starvation response of the bacteria *S. meliloti* which induces the production of 2 different daughter cells, one suited to short-term starvation and the other suited to longer-term starvation (Ratcliff and Denison, 2010). Also consistent with our results are recent laboratory evolution studies showing that (1) the frequency and duration of bet hedging (persistence) in bacteria is heritable (Rotem et al., 2010), (2) different patterns of antibiotic treatment can select for a high frequency of bet hedgers (Van den Bergh et al., 2016), (3) antibiotic tolerance is determined by evolution of cell cycle reentry timing (lag times) (Fridman et al., 2014), and (4) lag times evolve as a function of the duration of antibiotic treatment (Fridman et al., 2014). Together with our results these findings link bet hedging strategies in microorganisms with the diversity of bet hedging strategies in higher eukaryotes, drawing parallels for example with the pioneering studies of seed dormancy bet hedging in desert annuals (Cohen, 1967). In microorganisms, when cautious cells respond to environmental stress with longer lag times whose frequency can be tuned/evolve to any value between 0 and 1, we suggest this be called “tunable bet hedging” to distinguish it from previously described pre-adaptive “stochastic bet hedging”.

In yeast as in multicellular organisms, fitness depends on reproduction in capricious and potentially lethal environments whose severity and duration are also unpredictable. Similar to seeds in dormancy, microorganisms in nature spend a large fraction of their time in post-diauxic or quiescent phases where they are naturally stress resistant but must mitigate risks when they resume growth or reproduction in potentially lethal environments (Gray et al., 2004). While classical evolutionary models assign fitness directly to genotypes, mutations, and mean trait values without consideration of the genotype-to-phenotype map, molecular models provide detailed mechanisms of development but rarely consider the effects of natural genetic variation. Many studies of phenotypic diversity in yeast and bacteria are conducted in one or a few strains, but here we studied well-characterized osmotic stress signaling responses on a backdrop of natural variation in 50 yeast genotypes adapted to diverse ecologies. This enabled our identification of negative feedback controlling perfect adaptation and robust recovery of steady-state viability in exponential cultures experiencing moderate osmotic stress, and the combined strategies of predictive plasticity and diversified bet hedging in post diauxic cultures responding to more severe conditions. It is increasingly clear that labyrinthine developmental mechanisms – that are themselves controlled by genetic variation – translate genotypes into phenotypes with variable fidelity that can also be selected (Yvert et al., 2013). Evolution of the genotype-phenotype map across different environments enables the honing of predictive plasticity and selection on bi-stable states required for the evolution of bet hedging (Rotem et al., 2010; Rutherford, 2003).

## Experimental procedures

### Strain acquisition and deposition

Over 200 unique wild and industrial diploid strains of *Saccharomyces cerevisiae* were obtained from the fungal diversity collection of Centraalbureau voor Schimmelcultures (CBS), an institute of the Royal Netherlands Academy of Arts and Sciences in Utrecht, Netherlands (http://www.cbs.knaw.nl). Strains that were modified for this report are listed in Tables 1 and Table 1–table supplement S1. They have been deposited to the Yeast Genetic Resources Lab of the National BioResource Project in Osaka, Japan (http://yeast.lab.nig.ac.jp/nig/index_en.html/).

### Haploid *MAT* a library of wild and industrial genotypes

Our goal was make a large library of haploid derivatives of wild and industrial strains in which to survey the effects of genetic variation at the level of osmotic stress signaling and downstream response. A total of 50 strains have been completely validated, and many others are in the pipeline. Ours is the largest such library of which we are aware and the results we report here show it should be generally useful – e.g. we have a large and representative sample of natural variation with bet hedgers ranging from almost completely cautious to completely reckless and sufficient ability to detect statistically significant effects. The first step in our library construction pipeline was to delete the *HO* locus of each strain by replacement with the KanMX4 marker gene and “barcodes” to permanently label each strain while preventing homothalism (Table 1– table supplement S1)(Giaever et al., 2002; Shoemaker et al., 1996). The KanMX4 gene was PCR-amplified for this purpose with primers containing the barcode sequences(Wach et al., 1994). Next, kanamycin-resistant transformants were grown in pre-sporulation medium containing 10% glucose followed by sporulation under starvation conditions in 1% potassium acetate. Although the strains differ in their sporulation efficiency and optimal conditions (see strain information at http://www.cbs.knaw.nl), we found it was most efficient to put strains through repeated rounds of a general sporulation protocol rather than trying to optimize the conditions for each strain. The *MAT***a** haploids were identified by “schmoo” formation in 96-well plates containing alpha factor and confirmed by crossing to a G418-sensitive, clonNAT-resistant *MATalpha* tester strain and selection on double-antibiotic plates. Next we deleted the *URA3* gene using a standard gene deletion method and selected the *ura3*Δ clones by replica plating and selection on 5-FOA. Finally, *ho* and *ura3* deletions and the barcode sequences of each strain were verified by PCR and sequencing. Forty-nine wild strains and a laboratory strain meeting these criteria were used in this study (see Tables 1 and Table 1–table supplement S1 for strain details).

### Synthetic population of *GPD1*::GFP wild/lab diploids

The *MATalpha* laboratory strain BY4742 was transformed to create a stably integrated GPD1::*GFP* reporter (G01) using a deletion cassette containing a *URA3* marker for selection on SC-URA plates(Gietz and Woods, 2002; Wach et al., 1994). A synthetic “population” of diploids was created by mating each strain in the library of *MAT****a*** haploids (50 strains) with *MATalpha* G01 by mixing on SC-URA plates for 2 hours followed by streaking onto selective SC-URA+G418 plates. The 50 resulting wild/lab diploid strains all have 50% of their genes and the *GPD1*::GFP reporter from strain G01 in the BY4742 laboratory strain background (Table 1). After mating, it was necessary to screen for triploids or tetraploids, which express higher levels of *GPD1*::GFP and have higher tolerance to osmotic stress. Overnight cultures of wild/lab yeast were diluted 50-fold into fresh YPD+G418 and grown for an additional 4 hours, fixed by 1:3 dilution into cold ethanol and re-suspended in 20 ug/ml RNAse A to digest ribonucleic acids. Digested cells were stained with 30 ug/ml propidium iodide to label DNA and ploidy was determined by flow cytometry (FACS Calibur; Becton Dickinson).

### Exponential and post-diauxic cultures

Fresh cultures were generated for each experiment by replicating frozen 96-well plates onto YPD+G418 agar followed by 4 days growth at 21° C. To obtain mid-exponential cultures, freshly patched cells were grown in 2 ml liquid YPD+G418 cultures at with rotation (72 rpm) at 21° C. for 2 days. Two microliters of these suspensions were diluted into 2 ml of liquid YPD+G418 and grown at 21° C. for 14 hours (e.g. 5 rounds of cell division on average, with strain ODs ranging from 0.80 – 1.44). For post-diauxic cultures, freshly patched cells were grown in 2 ml liquid YPD+G418 cultures at with rotation (72 rpm) at 21° C. for 4 days. Strains cultured up to 8-days post-diauxic growth were tested for osmotic stress resistance and we found that 4 day cultures were already maximally resistant (not shown).

### Survival plating assays

To determine the adaptation limit of each strain (Table 3), post-diauxic cultures were diluted to OD_600_ of 0.1 with exhausted YPD (to prevent re-growth), sonicated for 5 seconds at a low setting (2.5; Sonifier Cell Disrupter, Model W185) and plated (5 ul) on 96-well YPD plates containing KCl ranging from 2.0 to 3.0 M. Growth was examined for up to 2 months at 21°C. Viability and static survival under osmotic stress (Figure 2B) was determined after incubation in 96-well microtiter plates containing liquid media with increasing concentrations of KCl for the times indicated, followed by plating on iso-osmolar YPD agar plates.

### Microfluidics

We used custom made microfluidics devices with two fluid inputs as described(Bennett et al., 2008). When performing microfuidics with post-diauxic cells, post-diauxic cultures were inoculated into devices with exhausted YPD medium and allowed to stabilize for a few hours prior to osmotic stress. Experiments were run at ambient room temperature and observed using a Nikon TS100 inverted microscope. Recordings were made using a Photometrics CoolSnap HQ2 digital camera operated by Metavue (Molecular Dynamics). Analysis of acquired images was performed using Image J software (https://imagej.nih.gov/ij/).

### Flow cytometry

For flow cytometry, after osmotic stress treatments 4 ml of PBS was added to each culture. Cells were isolated by centrifugation and resuspended in 1 ml PBS, transferred to FACS tubes, sonicated (5 seconds at level 3, Sonifier Cell Disrupter, Model W185) and stained with 3 ug/ml propidium iodide (PI) to monitor viability. After 20 min GFP fluorescence and viability were quantified using a FACS Calibur flow cytometer (Becton Dickinson) that had been calibrated prior to each use with SPHERO Rainbow Fluorescent Particles, 3.0 – 3.4 um (BD Biosciences). Flow cytometry data were gated using magnetic windows in FlowJo software to eliminate cell fragments, clumped and dead (PI-positive) cells (http://www.flowjo.com/).

For analysis, raw data for the viable cells in each sample (forward scatter, side scatter and GFP fluorescence data; up to 10,000 cells/sample) were extracted into an SQL database. Cell data were scaled for linearity (e.g. FLH1^1/3^, FSC^1/3^, SSC^1/2^ for GFP fluorescence, forward scatter, and side scatter, respectively). Distributions of *GPD1*::GFP accumulation in exponential cultures were unimodal, and therefore well-defined using a single mean (e.g. Figure 1–figure supplement S1). By contrast, *GPD1*::accumulations of cells in post-diauxic cultures were clearly multimodal at many time points (Figure 2–figure supplement 1). To identify different distributions of cells we used machine learning was performed using the sklearn.mixture option in the Gaussian Mixture Model (GMM) algorithm of the Python scikit package (http://scikit-learn.org/). The GMM algorithm identified parameters of the four most-likely Gaussian (defined by means and covariances) given the data for each sample. The 2-dimensional fits of *GPD1*::GFP and forward scatter data distinguished different cell types slightly better than fitting *GPD1*::GFP only; adding side scatter to fit distributions in 3-dimensional space little additional resolution. The number of Gaussians to be fit is a parameter that must be provided to the model. We used Bayesian information criteria (BIC) to determine that the data were well described by four distributions. In samples containing obviously fewer than four distributions, the under-populated distributions were assigned a correspondingly low frequency of cells.

### Clustering

To group, and ultimately rank, the strains according to their osmotic stress signaling responses to 2.5 M KCl during post-diauxic growth we used hierarchical clustering with Wards method in the fastcluster Python implementation (http://www.jstatsoft.org/v53/i09/)^34^. First we created state vectors of each strains behavior. Cell distributions were binned onto a 100 X 100 2-D grid according to their *GPD1*::GFP and forward scatter data, smoothed with Python scipyndimage.filters.gaussian_filter and normalized to define a linear 10,000 element state vector for each sample (strain, time point). While 100 bins on each axis where sufficient to capture detailed distributions while allowing efficient computation, we found stronger clustering when performing the same analysis using the combined *GPD1*::GFP and forward scattering data. The osmotic stress response up to 168 hours was defined by the vectors for each of the 7 time points, successively appended to form a 70,000 element time-line vector representing the combined evolution of *GPD1*::GFP accumulation and forward scatter data. The time-line vectors were used to compute a distance matrix between strains using the symmetric Kullback-Leibler divergence. As each strains and time point was replicated between 4 and 15 times, we controlled for variation in sampling and clustering outcomes by randomly drawing samples for each strain and time point with equal probability. Clustering was repeated for a total of 17,000 permutations requiring 43 hours of computation time on a 3.7 GHz Intel 7 iMac. This was sufficient to achieve stable Monte-Carlo statistics. Computational sorting of time-series distributions resolved 6 clades differentiated for rates of GFP accumulation, adaptation and survival. The fraction of permutations in which each strain grouped with more than half of the other strains in its mean cluster was used to rank that strain’s behavior relative to the other strains in its group (clustering statistics; Table 5).

## Author contributions

YH designed and performed experiments, data analysis, figures and writing for initial versions of this manuscript. SB performed modeling, database construction, data analysis, statistical design, computer programming, writing and editorial support. WLP designed and trained us in microfluidics devices. SR conceptual design, workflow for wild strain collection, data analysis, model development, figures and writing.

## Acknowledgments

We thank Lisa Castenada, Tanya Gottlieb, Adrianne Hughes, Brandon (Tommy) Huynh, Katie Michaelsen, Rachel Rodman, and numerous undergraduate helpers for construction and validation of the wild strain collection, Jeff Hasty for advice on microfluidics and Stéphane Douady and Pierre-Yves Bourguignon for discussions and advice on scaling and clustering. This research was supported by a Scholar Award from the Damon Runyon Cancer Research Foundation (SR) and the Lady Tata Memorial Trust (YH).

## Tables

**Table 1– table supplement S1.**
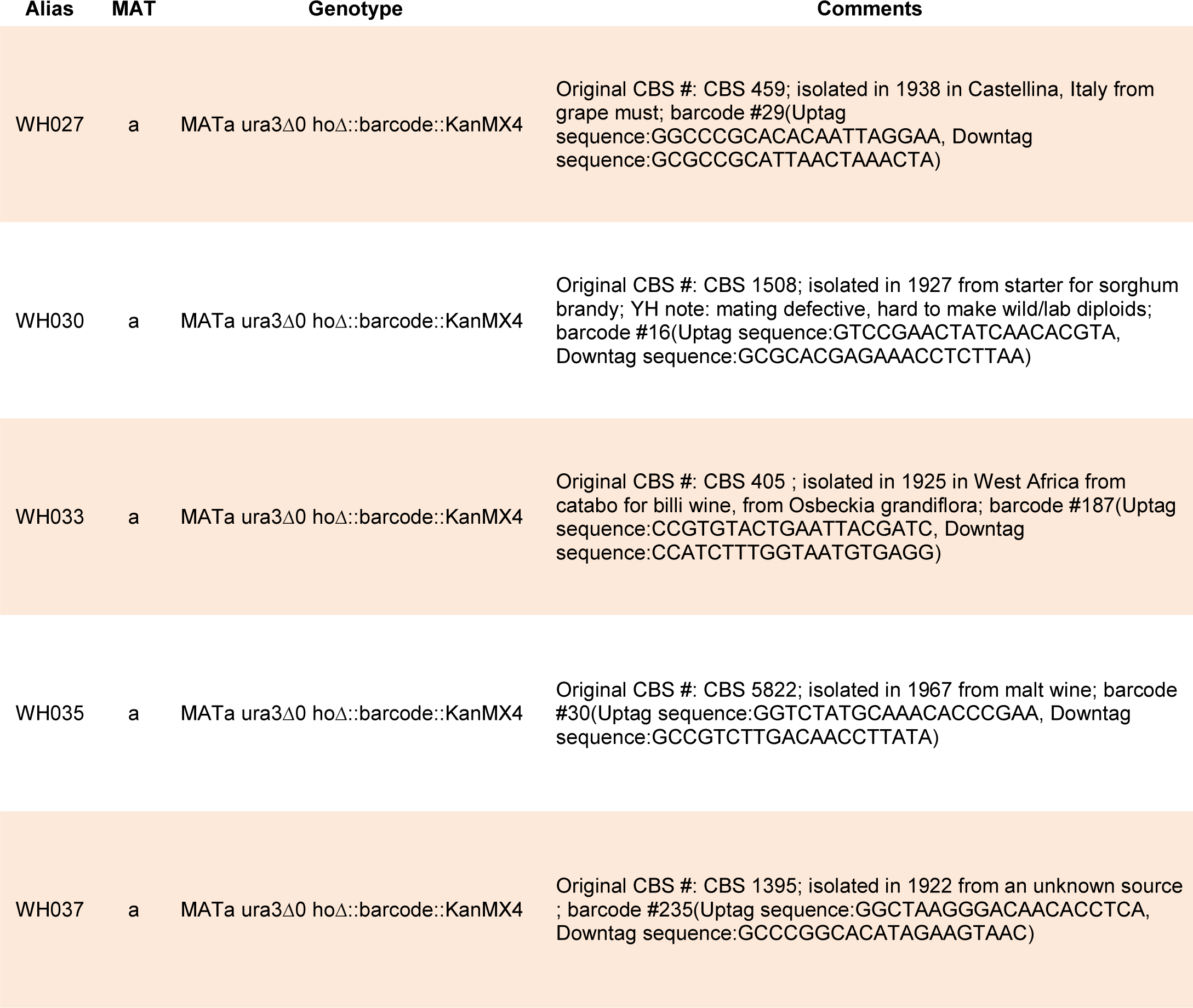
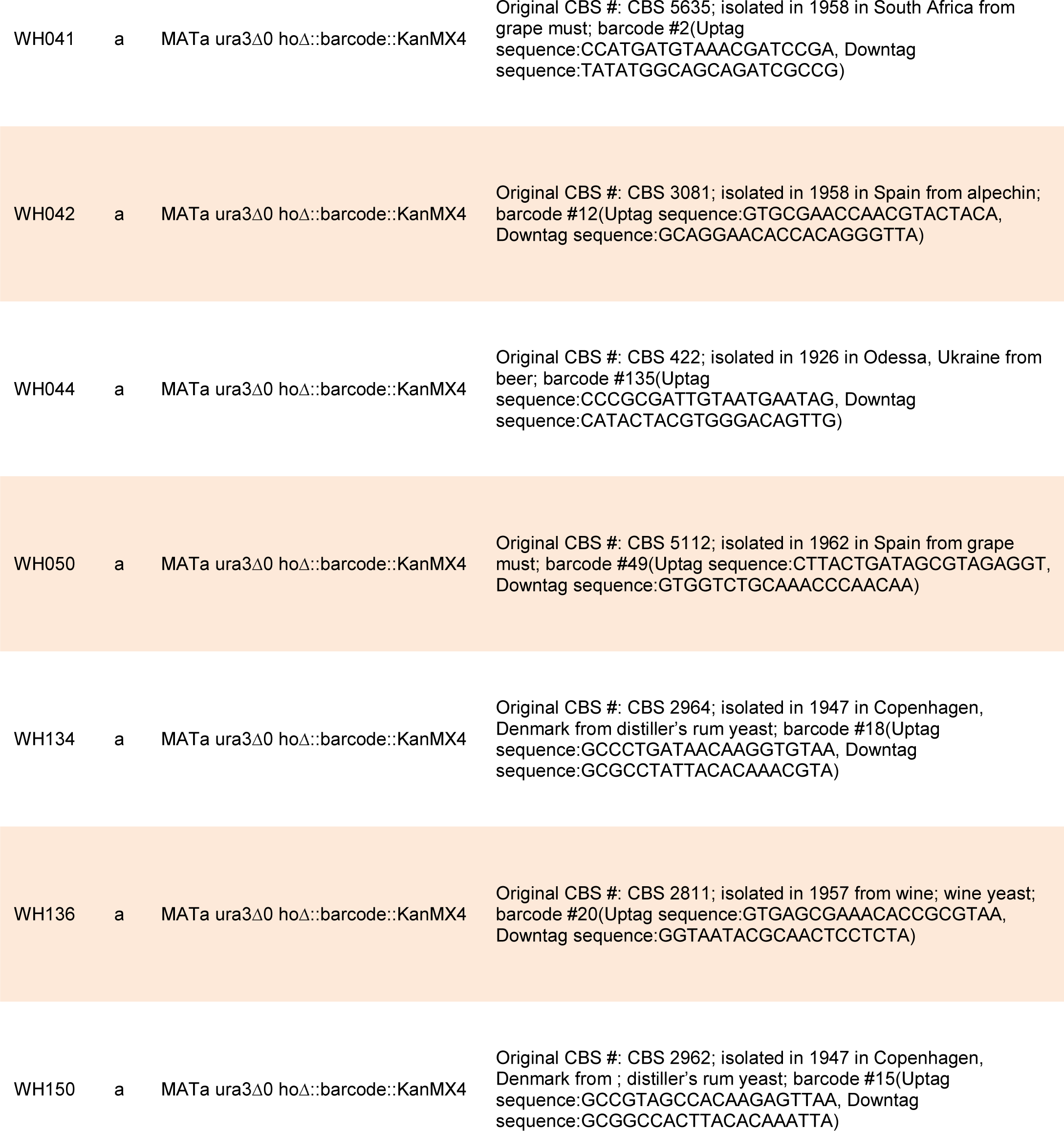
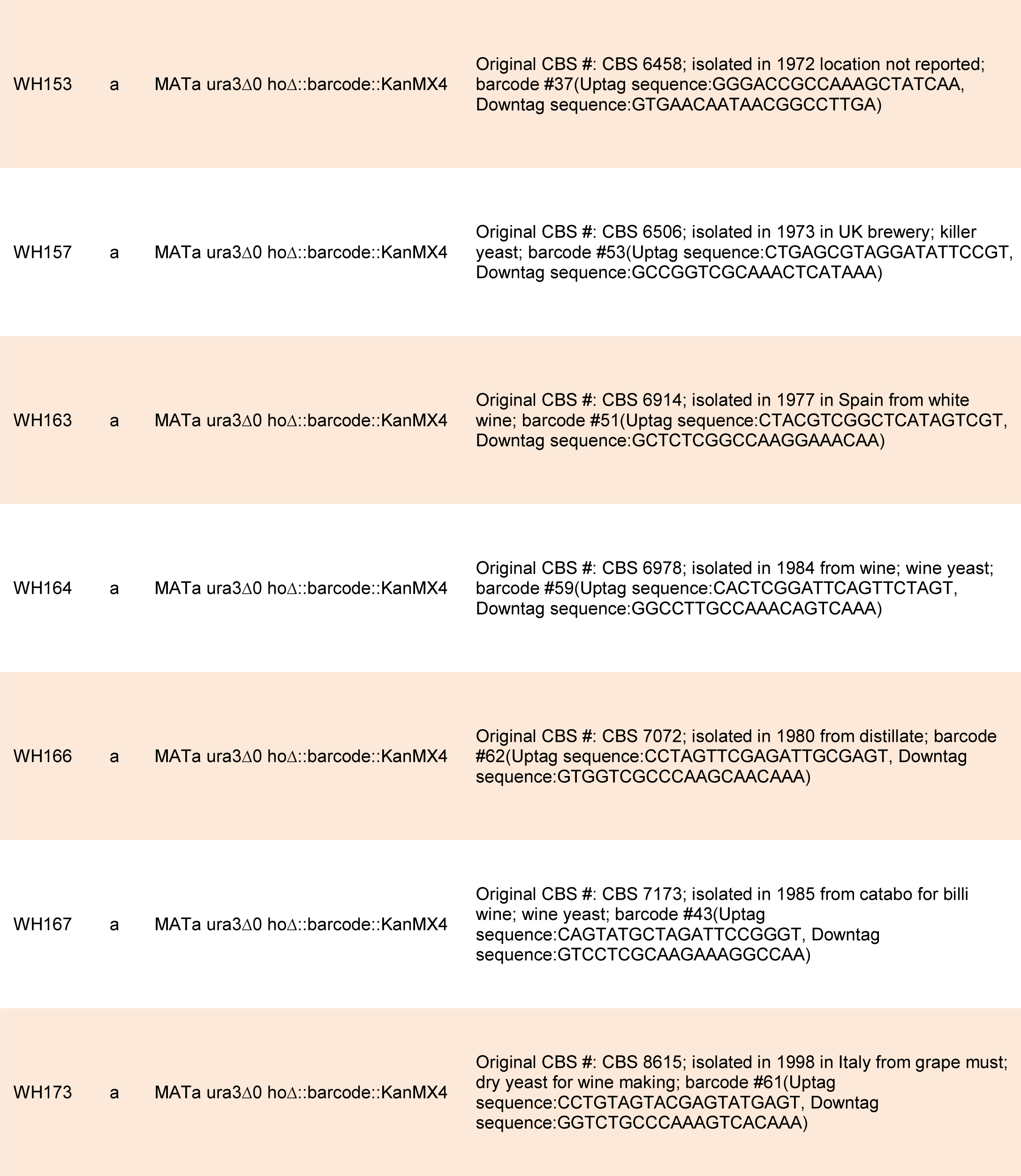
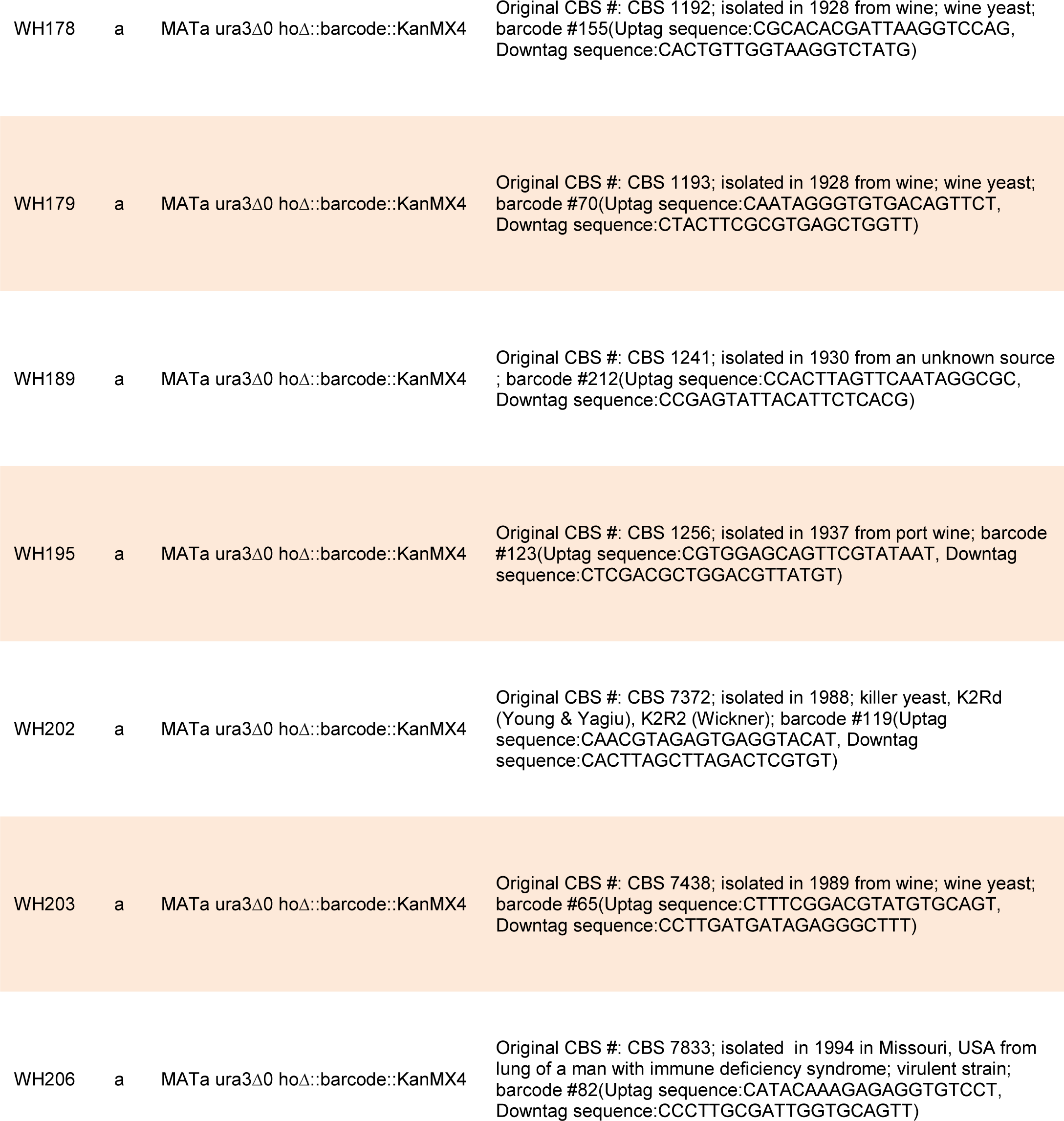
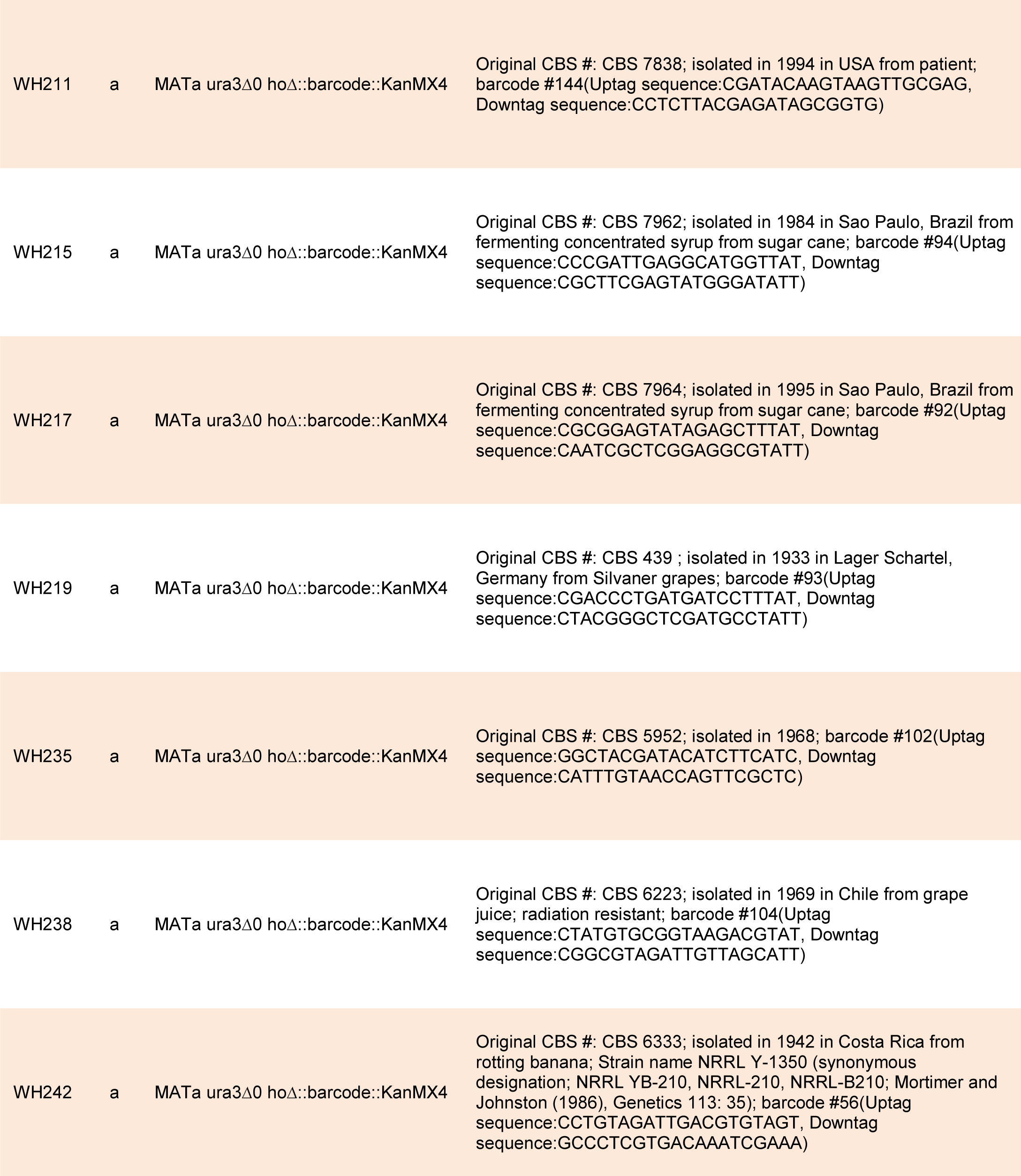
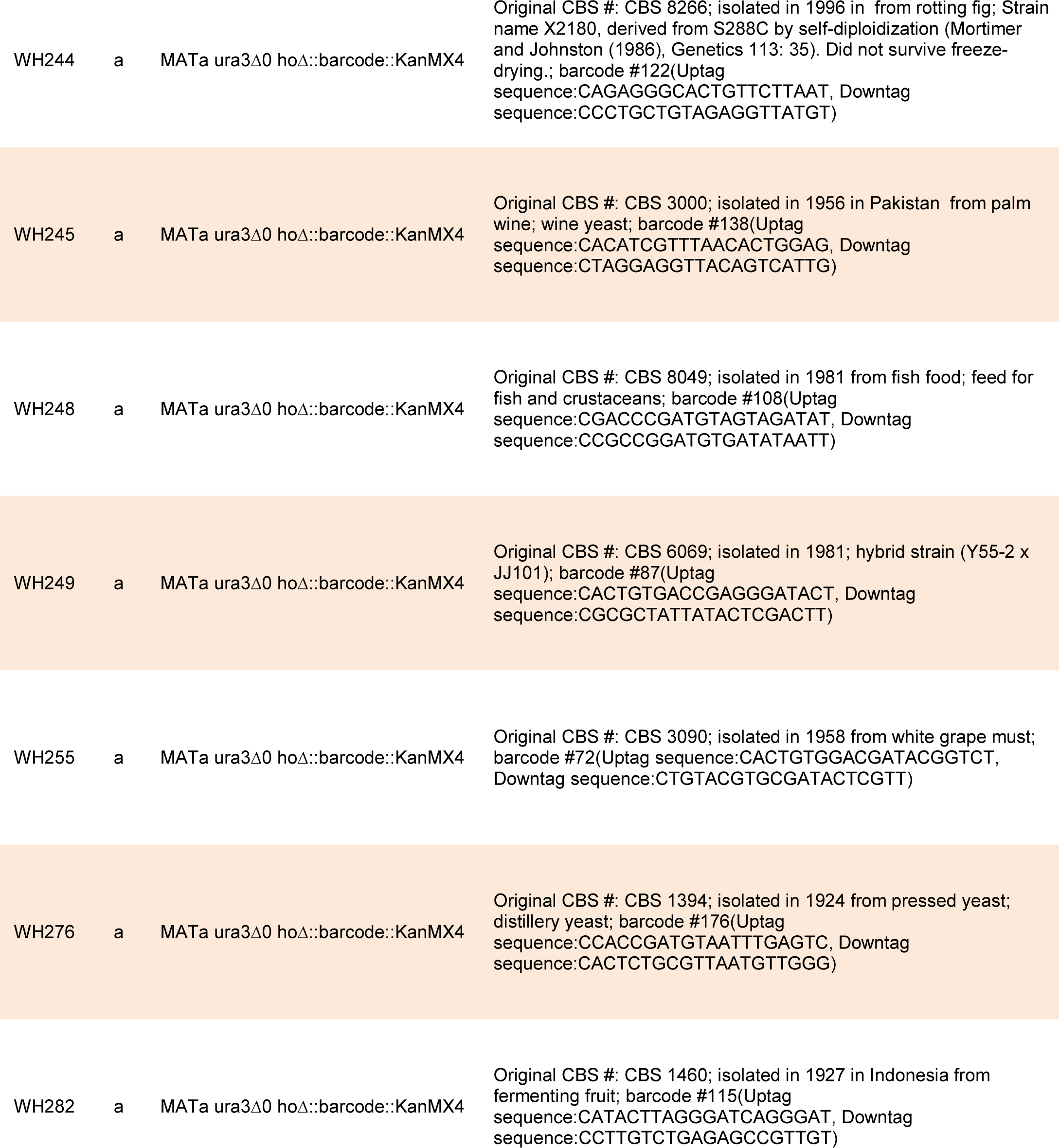
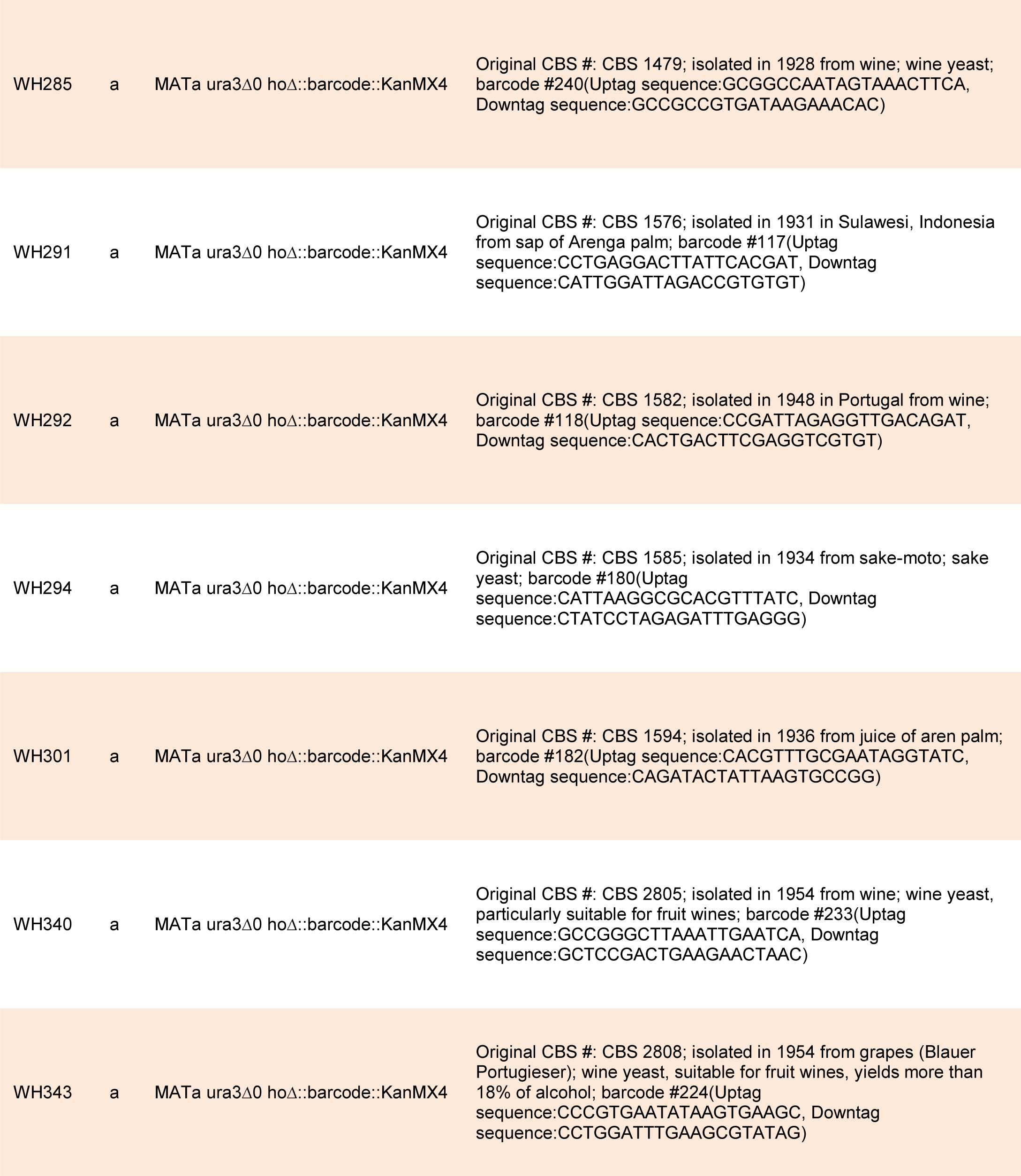
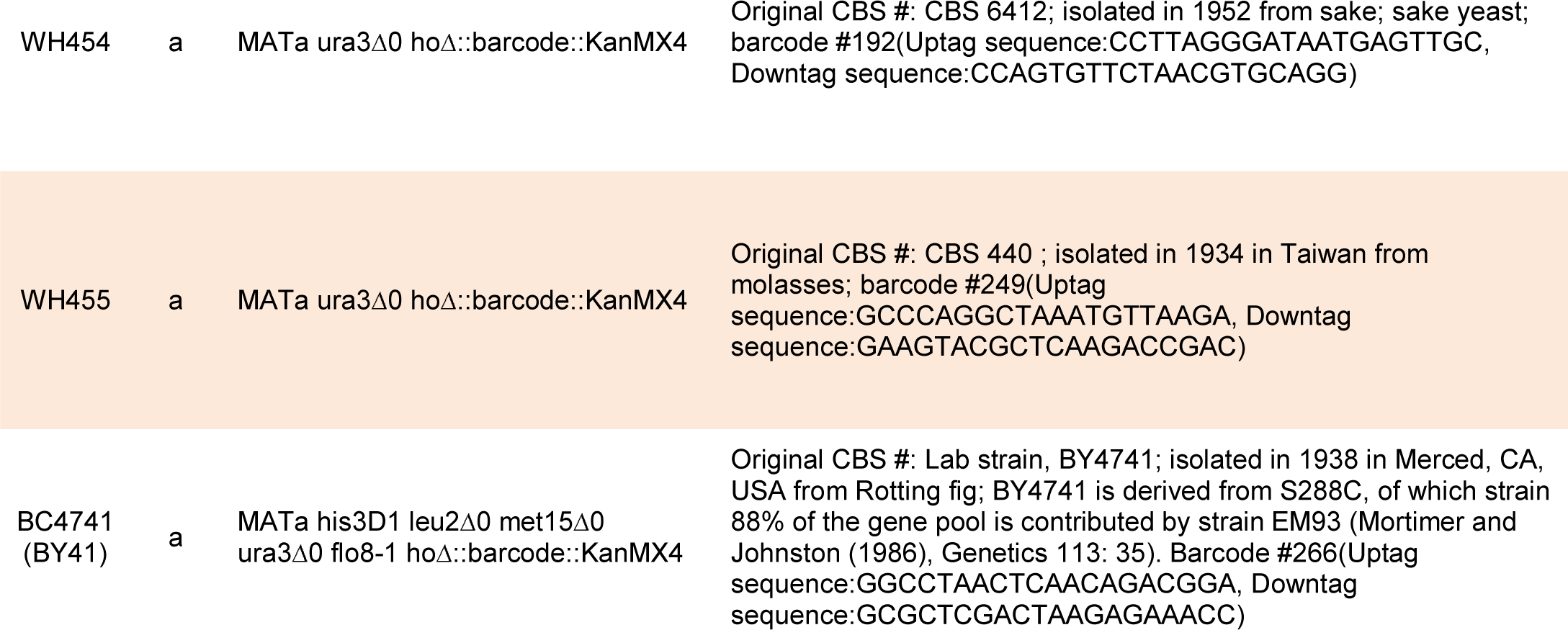
Haploid derivatives of wild strains. The source for all wild strains in this study was the strain collection of the Royal Netherlands Academy of Arts and Sciences over the past 100 years (Table 1 and Table S1). All strains used in this study have been deposited to the Yeast Genetic Resources Lab of the National BioResource Project in Osaka, Japan http://yeast.lab.nig.ac.jp/nig/index_en.html.

## Figures

(High resolution figures available at https://figshare.com/s/222fd592f52f59d5d4fb)

**Figure 1–figure supplement 1.**
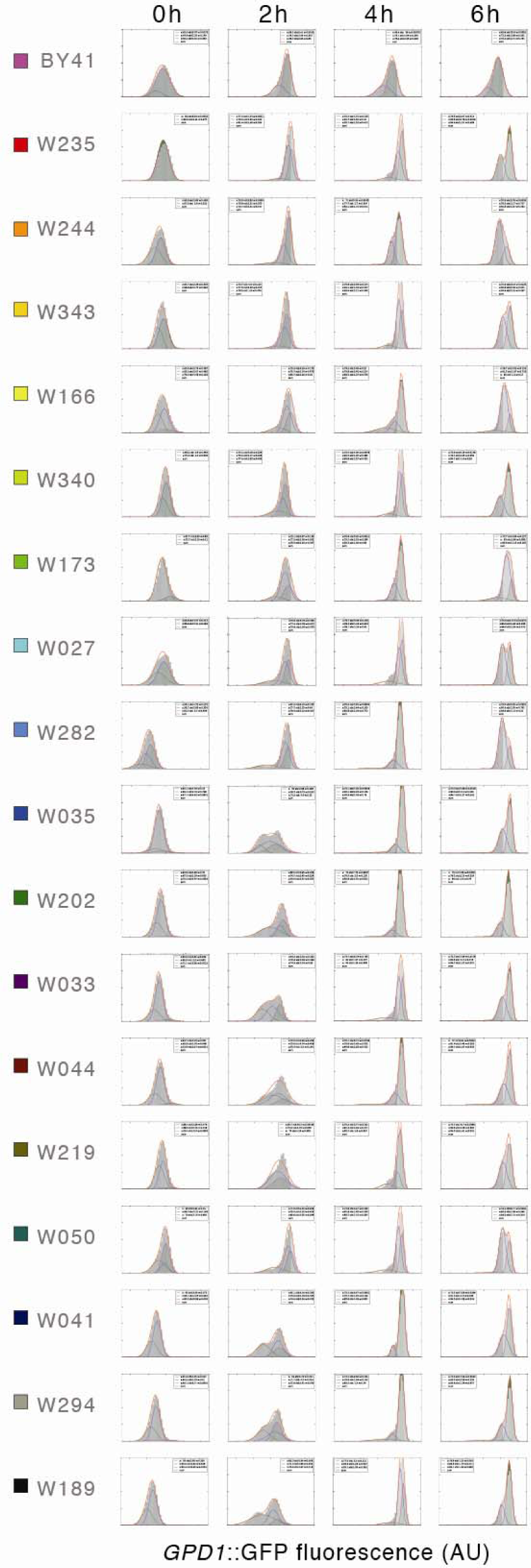
Mean *GPD1*::GFP accumulations accurately track the rate of osmotic stress signaling in exponential cultures. Representative distributions of cells from exponential cultures exposed to 0.75 M KCl for the times shown were generally monomodal and well-approximated by mean values. Learned frequency distributions of *GPD1*::GFP accumulation (AU) with mean (x), standard deviation (std), and weight (w; the fraction of cells in each distribution) shown in boxes (zero-weighted distributions not shown). Sum (red) shows the cumulative fit of the 4 learned Gaussians. *GPD1*::GFP values were normalized across all strains for comparison. The 18 representative strains are color-coded as in Figure 1B.

**Figure 2–figure supplement 1.**
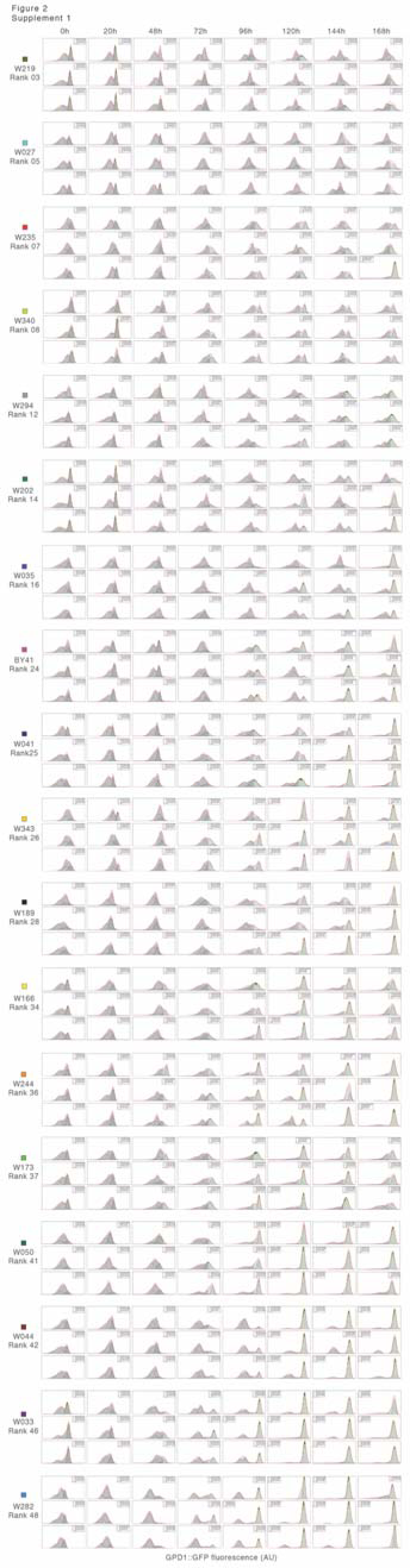
Reproducible rates of *GPD1*::GFP accumulation in post diauxic cultures during extreme hyperosmotic stress. Representative replicates of learned distributions of GPD1::GFP accumulation in post-diauxic cultures exposed to 2.5 M KCl for the times shown. Mean (x), standard deviation (std), and weight (w; the fraction of cells in each distribution) are given (zero-weighted distributions not shown). Sum (red) shows the cumulative fit of the 4 learned Gaussians. The 18 strains shown are color-coded as in Figure 1B.

**Figure 2-figure supplement 2.**
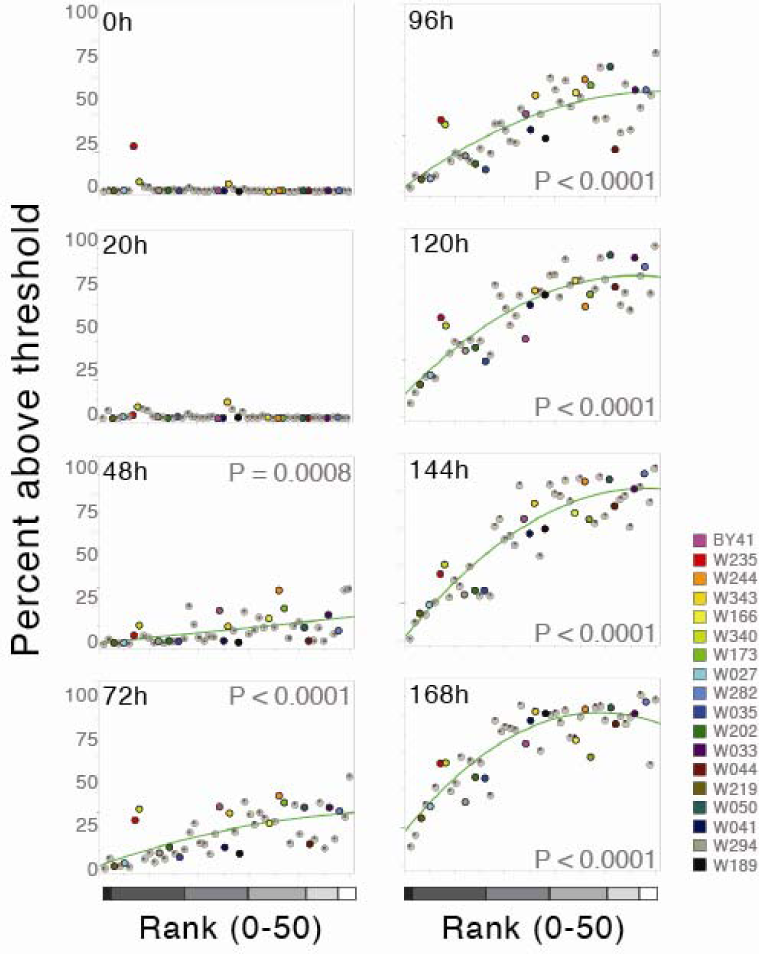
Rank predicts rate of *GPD1*::GFP accumulation. Strains were exposed to 2.5 M KCl for increasing times shown and ordered according to rank. The average percent of cells in each strain above a threshold set at the top 11% of accumulation of *GPD1*::GFP normalized across all post-diauxic cultures. Significant P-values for 2^nd^ order quadratic fits of the data are shown (JMP, SAS Institute; N=50 strains). The number of strains in each mean cluster is indicated with increasingly lighter grey scale in their order of ‘cautious’ to ‘reckless’ signaling: 2 strains (MC0), 15 (MC1), 12 (MC3), 12 (MC2), 6 (MC5), 3 (MC4). The 18 representative strains are color-coded as in Figure 1B.

**Figure 2–figure supplement 3.**
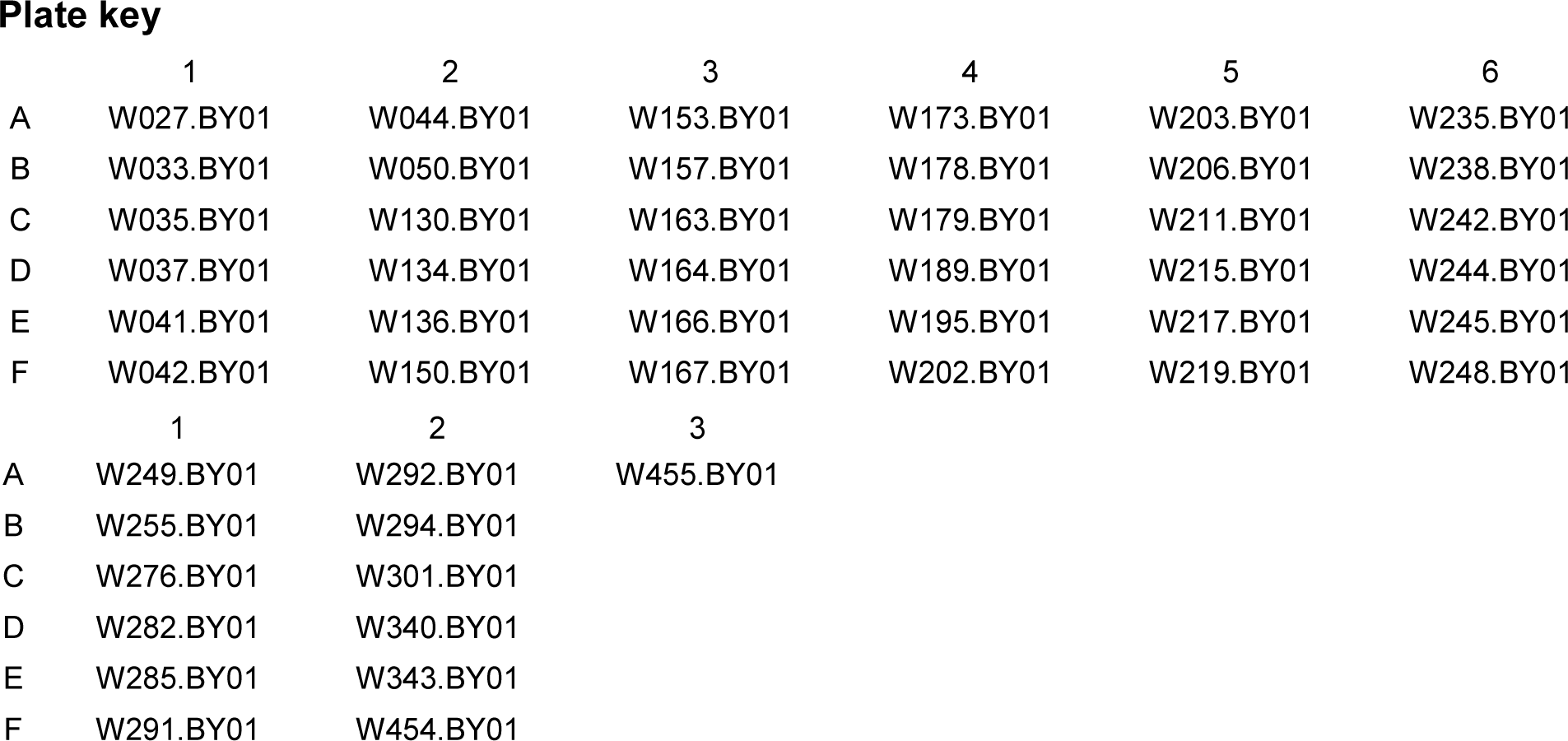
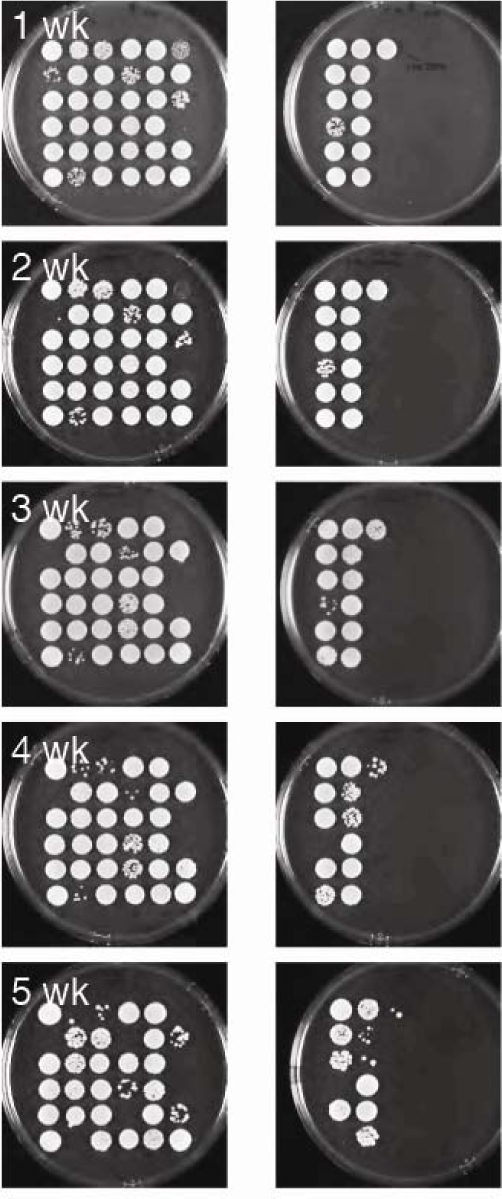
Regrowth after static viability and survival in extreme hyperosmotic stress. Raw data used for montage for Figure 2B shown as plated. Post-diauxic cultures were incubated for up to 5 weeks in 3 M KCl and then plated on iso-osmolar media.

## Movie Legends

**Movie 1.** Exponential W027 cells seeded with a single post diauxic cell of the same genotype (box). Media was switched to 1.5M KCl at time 0, GPD1::GFP fluorescence is shown in green. Time stamp shown in upper right.

**Movie 2.** Post-diauxic W027 cells exposed to 1.5 M KCl, GPD1::GFP fluorescence is shown in green. Time stamp shown in upper right.

**Movie 3.** Post-diauxic W027 cells exposed to 2.5 M KCl, GPD1::GFP fluorescence is shown in green. Time stamp shown in upper right.

## Bet hedging model

Annotated code for our model of bet hedging with heritable probability of binary, cautious versus reckless bet hedging is publicly available (https://figshare.com/s/2c03544aef0c40cc86c2). The bet hedging ‘strategy’ P was defined as the heritable probability of cautious cells for 0 ≤ P ≤ 1. Nine possible 2-state environmental shifts between three general osmotic stress environments were considered: permissive (E0; all cells grow equally well), restrictive (E1; reckless cells divide, cautious cells survive without dividing), and killing (E2; reckless cells die, cautious cells survive without cell division). The relative fitness of representative strategies (0, 0.1, 0.2,…1.0; number of surviving cells in each strategy divided by the total number of surviving cells across all strategies) was calculated after 10 generations in each environment except as shown on Figure 3c. For simplicity, the natural attrition of older cells (death and disappearance) and rates of cell division were assumed to be equal for all strains. Results were independent of the number of generations in the first environment except as shown when E1 was the first environment.

## Databases and linked archives

Flow cytometry database (annotated) https://figshare.com/s/52ef966b16cba7f41d7f
Python script for bet hedging model https://figshare.com/s/2c03544aef0c40cc86c2
Figure 1–figure supplement S1 complete data set https://figshare.com/s/8b709fd16cccbabc2a5a
Figure 3–figure supplement S3 complete data set https://figshare.com/s/8147275b62eb8d4db6bf
Excel file with tables and raw data https://figshare.com/s/00a7bf31d2791922f1d8

